# Implicit Hierarchical Tensor Decomposition of Single-Cell Four-Omics Data Reveals Cell-Type-Linked Candidate Enhancer–Promoter-Associated Programs

**DOI:** 10.64898/2026.08.18.745447

**Authors:** Y-H Taguchi, Turki Turki

**Affiliations:** Department of Physics, Chuo University, 1-13-27 Kasuga, Bunkyo-ku, Tokyo 112-8551, Japan; Department of Computer Science, King Abdulaziz University, Jeddah 21589, Saudi Arabia

**Keywords:** single-cell multi-omics, tensor decomposition, HOSVD, enhancer–promoter regulation, chromatin accessibility, H3K27ac, 3D genome, gene regulation, CHARM

## Abstract

Single-cell multi-omics provides complementary views of gene regulation, but integrating modalities with natural features comprising enhancers, genes, and three-dimensional (3D) genomic loci remains challenging. This study developed an implicit hierarchical tensor decomposition framework and applied it to the CHARM single-cell four-omics mouse brain data (GSE303006) by jointly analyzing ATAC, H3K27ac, RNA, and reconstructed 3D enhancer–promoter (E–P) distances. After quality control, 730,969 E–P pairs, 42,669 enhancers, 16,239 genes, 391,435 20-kb bin pairs, and 4,258 cells were retained. Each modality was independently reduced to 20 components and mapped to an implicit **730,969 *×* 4,258 *×* 20 *×* 4** tensor, which was decomposed without materializing the full array. In a joint model, cell type explained more variance than biological replicate for all 12 distinct cell components represented among the 100 largest Tucker core elements. A representative core linked the Inh_Ndnf/Lamp5–Ex_L3/4_IT cell axis to contrasting presynaptic-transmission and developmental/morphogenetic candidate E–P-associated programs. Its E–P and cell factors were recovered when the whole analysis was repeated independently on each pair of replicates, and the E–P pairs it prioritized agreed in direction with the cell-type-resolved enhancer– gene links of the original CHARM study. No single modality reproduced the program, which required the combination of accessibility, chromatin contact, and expression. Backprojection showed reproducible ATAC, H3K27ac, and 3D effects across three biological replicates; RNA differed clearly for the developmental module but not for the presynaptic module. The framework thus identifies regulatory structure that is shared across molecular modalities and is not recoverable from any single modality.

## 1 Introduction

Cellular identity is established through multiple regulatory layers, including chromatin accessibility, histone modifications, three-dimensional (3D) genome organization, and transcription. Recent single-cell technologies have begun to measure several of these layers simultaneously, enabling direct investigation of the coupling between regulatory state and transcription in heterogeneous tissues [1–3]. The CHARM method is particularly relevant because it profiles genome conformation, histone modifications, chromatin accessibility, and RNA in the same single cells and has been applied to mouse embryonic stem cells and cortical tissues [3]. The resulting dataset therefore provides an unusual opportunity to investigate regulatory relationships at single-cell resolution without requiring cross-assay cell matching.

Most multimodal integration frameworks are primarily cell-centric, meaning that they seek a common cell embedding or shared neighborhood structure across modalities [4, 5]. However, for enhancer–promoter (E–P) regulation, the feature spaces are intrinsically heterogeneous. ATAC and H3K27ac are naturally enhancer-level signals, RNA is gene-level, and 3D organization is defined for pairs of genomic loci. Direct concatenation can obscure distinct feature identities, whereas expanding every modality to all candidate E–P pairs creates an extremely large and highly redundant representation.

Biologically, distal enhancers regulate gene expression by coming into spatial proximity with their target promoters, and the four CHARM measurements describe different parts of this relationship: accessibility and H3K27ac at the enhancer, expression of the gene, and the 3D distance between the two. The candidate E–P pair is therefore the smallest unit at which all four measurements are defined. Using it as the common feature axis allows the four modalities to be placed in a single tensor, and it makes the result interpretable in regulatory terms, because an E–P factor ranks enhancer–promoter relationships rather than the features of a single assay. The pairs considered here are candidates (all TSSs within 1 Mb of an enhancer), not established functional links.

Tensor methods provide a natural framework for multimodal data because they retain multiple modes rather than flattening them into a single feature matrix. Higher-order singular value decomposition (HOSVD) generalizes matrix SVD to multiway arrays and yields orthogonal factors for each mode together with a core tensor that describes their interactions [6]. However, directly constructing a tensor containing hundreds of thousands of E–P pairs, thousands of cells, multiple latent components, and several modalities is computationally impractical.

This study develops a hierarchical strategy that avoids this materialization. Each modality is first decomposed independently into 20 components. The resulting modality-specific feature scores are then mapped to E–P pairs with inverse-square-root degree correction, while the original stage-1 component index is retained as an explicit tensor mode. The second-stage HOSVD is computed from compact Gram matrices and blockwise projections. This produces an interpretable E–P factor, cell factor, component mode, and modality factor for a conceptual tensor of dimensions 730,969×4,258×20×4. The procedure is summarized in Figure 1, and its full derivation is given in Supplementary Note 1.

**Fig. 1.**
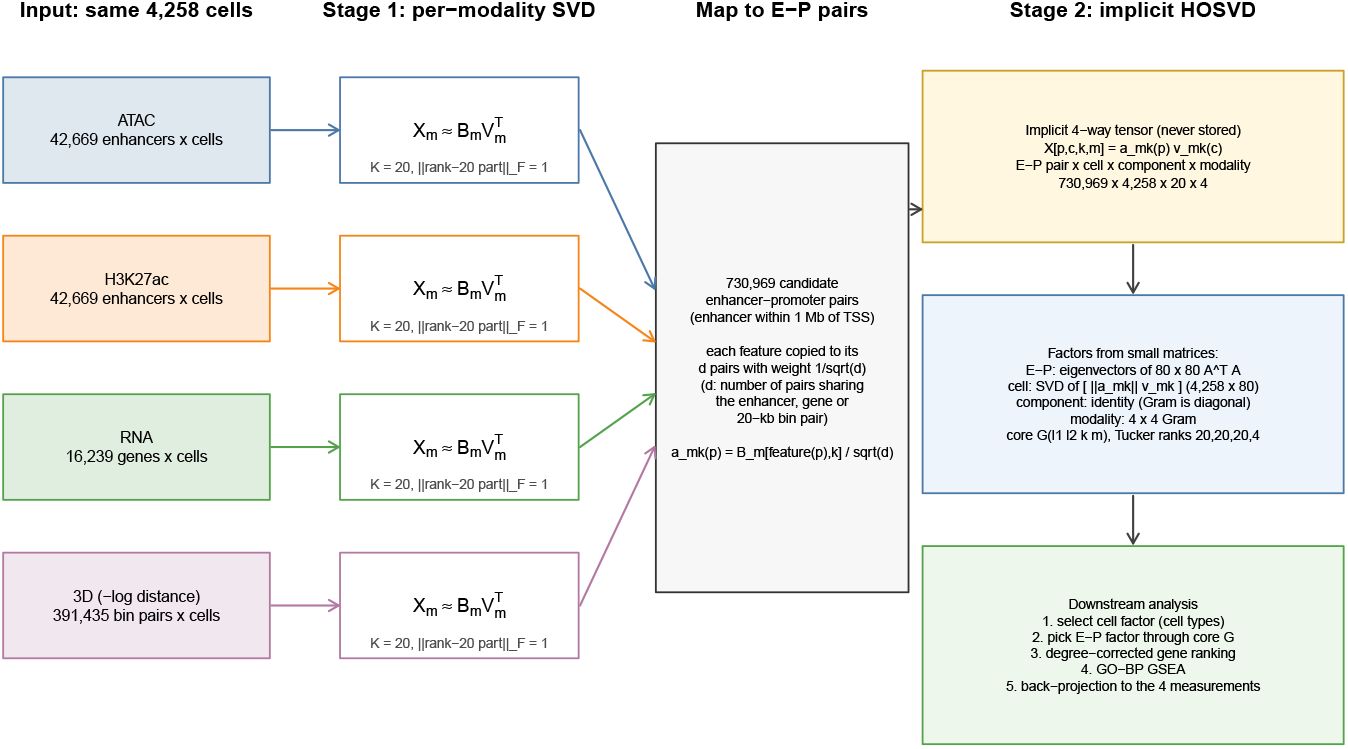
Overview of the implicit hierarchical tensor decomposition. Each of the four modalities measured in the same 4,258 cells is decomposed independently by a rank-20 SVD (stage 1). The feature scores are copied to the 730,969 candidate enhancer–promoter pairs with inverse-square-root degree weights, which defines an implicit E–P pair *×* cell *×* component *×* modality tensor that is never stored. Its HOSVD (stage 2) is obtained from small matrices: an 80 *×* 80 Gram matrix for the E–P mode, a 4,258 *×* 80 matrix for the cell mode, an identity component factor, and a 4 *×* 4 Gram matrix for the modality mode. Downstream, a cell factor associated with cell types is selected, the E–P factor coupled to it through the core tensor is converted into a degree-corrected gene ranking, and the result is examined by GO enrichment and by backprojection to the original measurements.

We applied this framework to the mouse brain CHARM dataset GSE303006 [3]. This study aimed to (i) determine whether the dominant cell-mode structure reflects biological cell type rather than replicate-specific variation; (ii) assess whether individual large Tucker-core elements can be mapped back to coherent candidate E–P-associated programs; (iii) determine whether those latent programs are visible directly in the original ATAC, H3K27ac, RNA, and 3D measurements; and (iv) test whether these programs are robust to the replicate structure and to the choice of modalities, and whether the prioritized pairs agree with independently inferred enhancer–gene links. The results identify a general cell-type-dominated structure among the largest core elements and a representative regulatory axis linking neuronal subtype identity to distinct synaptic and developmental candidate E–P-associated programs.

## 2 Results

### 2.1 Quality-Controlled Four-Omics E–P Representation

The initial E–P construction paired active enhancer candidates with expressed genes within 1 Mb. Following RNA, enhancer, 3D, and cell quality control, the final analysis contained 730,969 unique enhancer–gene pairs. The retained feature counts are summarized in Table 1.

**Table 1.**
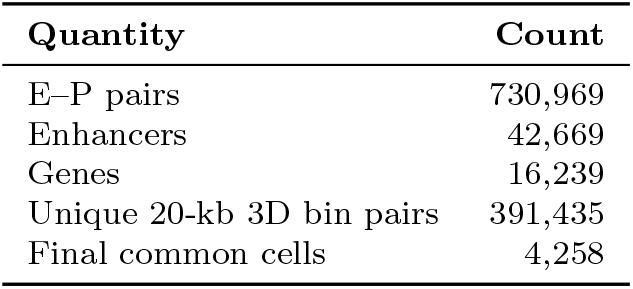
Final dimensions of the four-omics E–P analysis.

| **Quantity** | **Count** |
|---|---|
| E–P pairs | 730,969 |
| Enhancers | 42,669 |
| Genes | 16,239 |
| Unique 20-kb 3D bin pairs | 391,435 |
| Final common cells | 4,258 |

Before enhancer detection filtering, 1,032,664 E–P pairs passed the 3D criterion. Requiring both enhancer ATAC and H3K27ac detection fractions to be at least 0.005 reduced the set to 730,969 E–P pairs. Enhancer degree *d_e_* ranged from 1 to 76 (median 15), gene degree *d_g_* from 1 to 174 (median 43), and 20-kb bin-pair degree *d_b_* from 1 to 36 (median 1). The inverse-square-root degree identities were verified numerically.

### 2.2 Hierarchical Decomposition Captures a Compact Multimodal Structure

Stage 1 produced feature-score matrices of 42,669 × 20 for ATAC, 42,669 × 20 for H3K27ac, 16,239 × 20 for RNA, and 391,435 × 20 for 3D. The corresponding cell-loading matrices were numerically orthonormal, with orthogonality errors on the order of 10*^−^*^14^. The rank-20 truncation retained 1.44%, 1.00%, 6.50%, and 9.31% of the variance of the standardized ATAC, H3K27ac, RNA, and 3D matrices, respectively (relative reconstruction errors 0.993, 0.995, 0.967, and 0.952). The cumulative retention for *k* = 1–20 and the reconstruction errors at both stages are given in Supplementary Table S1.

The second-stage conceptual tensor had 730,969×4,258×20×4 entries but was processed implicitly. The E–P mode had a numerical rank of at most 80, reflecting the four modalities times 20 stage-1 components. The top 20 E–P directions captured 63.03% of the total tensor energy when considered as an E–P-mode projection, and the top 20 cell directions captured 75.77%. The final 20×20×20×4 Tucker core retained 58.41% of the conceptual tensor Frobenius energy (relative reconstruction error 0.645). By modality, the core captured 48.9% of the ATAC, 33.0% of the H3K27ac, 67.5% of the RNA, and 84.2% of the 3D part of the conceptual tensor. Because the degree weights preserve norms exactly (deviation below 10*^−^*^14^), these fractions refer to the rank-20 stage-1 reconstructions; relative to the standardized original matrices, the overall retained fractions are the products of the two stages (0.71%, 0.33%, 4.39%, and 7.84%).

The modality-mode Gram matrix exhibited the largest off-diagonal similarity between ATAC and H3K27ac (0.161), whereas similarities involving RNA or 3D were substantially smaller. Thus, the latent tensor did not simply collapse all four modalities into a single common signal.

### 2.3 Cell-Type Effects Dominate Replicate Effects across the Largest Core Elements

The 100 largest absolute Tucker-core elements involved 12 distinct cell components. Every one of these 12 components had a larger cell-type Kruskal–Wallis effect size than replicate effect size (Figure 2). Thus, 100/100 top-core entries used a cell component lying above the 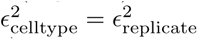 diagonal (a descriptive sign test is given in Supplementary Table S4).

**Fig. 2.**
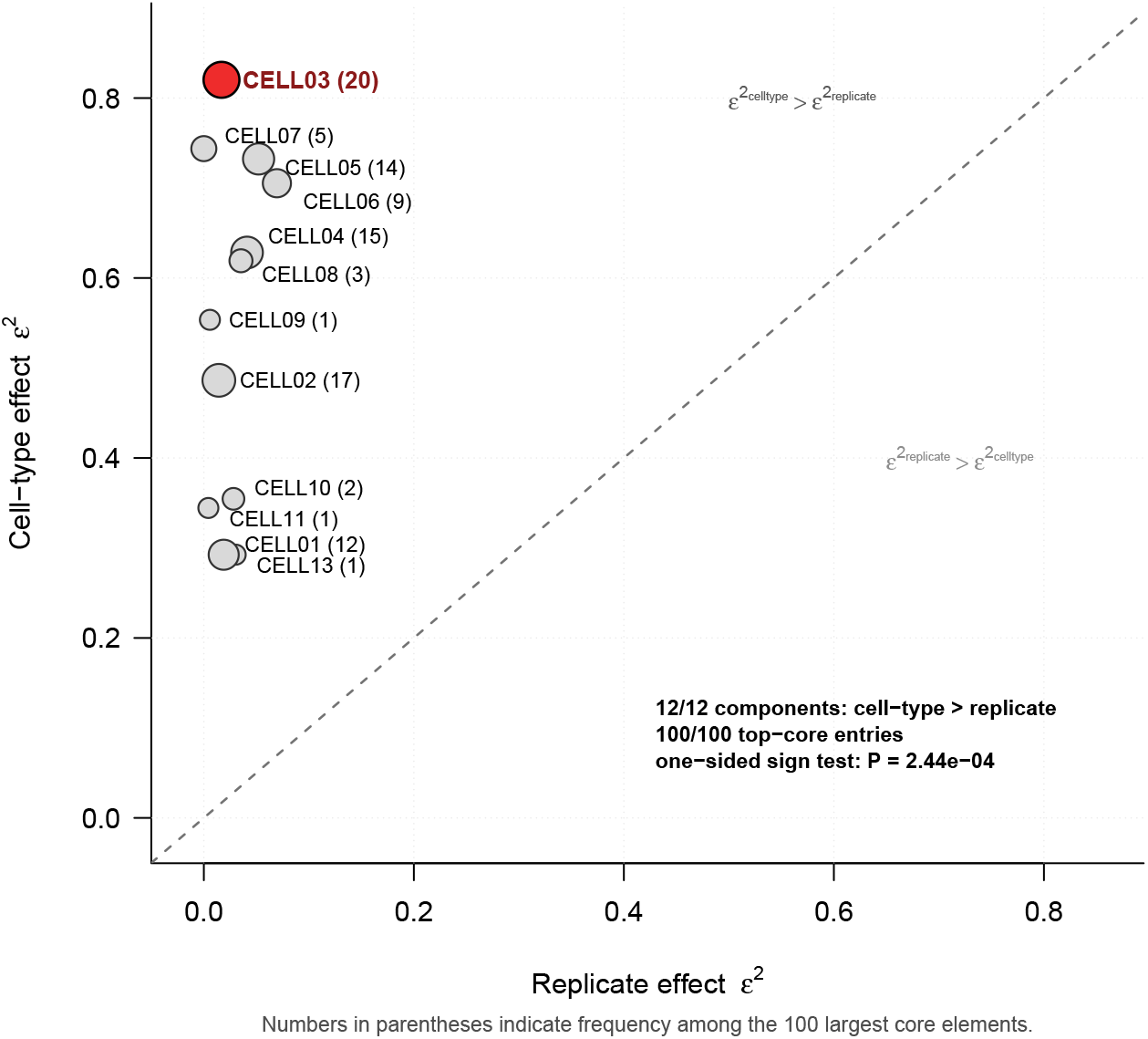
Cell-type effects dominate replicate effects across cell components represented among the 100 largest Tucker core elements. Each point represents one distinct cell-mode component. The horizontal axis shows the Kruskal–Wallis effect size (*ɛ*^2^) for biological replicate and the vertical axis the corresponding effect size for annotated cell type. Numbers in parentheses indicate how many of the 100 largest core elements contain each component. All 12 distinct components lie above the equality line; therefore all 100 top-core entries use a cell component with a larger cell-type than replicate effect. CELL03 is highlighted because it was selected for detailed backprojection.

Median *ɛ*^2^ across the 12 components was 0.586 for cell type and 0.0236 for replicate. CELL03, which was used in 20 of the 100 largest core elements, had the strongest cell-type effect: 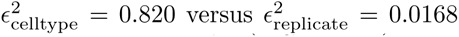. Other examples included CELL07 (0.744 versus approximately 0), CELL05 (0.732 versus 0.052), and CELL06 (0.705 versus 0.070).

These marginal comparisons do not by themselves separate the two factors, because cell-type composition differed between replicates (*χ*^2^ = 812, *P* = 1.4×10*^−^*^144^, Cramér’s *V* = 0.31; Supplementary Table S4). We therefore fitted, for each of the 12 components, a joint linear model with cell type, replicate, their interaction, and sequencing-depth and quality covariates. Cell type explained more variance than replicate and than the interaction for all 12 components (median type-II partial *η*^2^: 0.71 for cell type, 0.015 for replicate, and 0.034 for the interaction). The interaction explained more variance than the replicate main effect (median partial *η*^2^ 0.034 versus 0.015), indicating that replicate differences are not uniform across cell types. Within-cell-type tests confirmed this: across the 12 components and the 18 cell types with at least 20 cells in at least two replicates, replicate differences were significant (Benjamini– Hochberg-adjusted *P <* 0.05) in 77 of 216 tests. Most of these effects were small (median *ɛ*^2^ per component 0–0.07), but some were larger, most often in Inh_MSN cells (maximum *ɛ*^2^ = 0.44; Supplementary Table S4).

For CELL03, the interaction reflected a replicate shift confined to one of the two contrasted cell types. Ndnf/Lamp5 cells had lower CELL03 loadings in R2 than in R1 and R3 (rank-biserial *r*_rb_ = 0.69 and *−*0.63; Benjamini–Hochberg-adjusted *P ≤* 6 *×* 10*^−^*^5^; within-cell-type *ɛ*^2^ = 0.29), whereas L3/4 IT cells showed no replicate difference (|*r*_rb_|≤0.06). The largest replicate shift was 14.7% of the difference between the two cell types for Ndnf/Lamp5 and 0.24% for L3/4 IT, and the two cell types were completely separated by CELL03 within each of R1, R2, and R3 (rank-biserial *r*_rb_ = 1.00). The contrast that defines core 12 is therefore preserved in every replicate.

The cell-type structure of all 12 components is summarized in Supplementary Figure S3. Most components contrasted glial classes (astrocytes, the oligodendrocyte lineage, and microglia) with neurons, or neuronal subclasses with one another. CELL03 separated inhibitory interneurons (Pvalb, Sst, Ndnf/Lamp5, and Vip) from L3/4 IT excitatory neurons.

### 2.4 Core 12 Links a Neuronal Cell Axis to an E–P Regulatory Axis

Core rank 12 was the element *G_abkr_* with (*a, b, k, r*) = (3, 3, 2, 1), that is, E–P factor 3 (EP03), cell factor 3 (CELL03), stage-1 component 2, and modality factor 1 (MOD01), with value 0.168879. Its cell component was CELL03. Across all 4,258 cells, CELL03 was strongly structured by annotated cell type. Among the 50 largest positive loadings, Inh_Ndnf/Lamp5 cells were enriched; among the 50 most negative loadings, Ex_L3/4_IT cells were enriched. Consequently, CELL03 defined a strong inhibitory-versus-excitatory neuronal cell-state contrast while remaining comparatively insensitive to replicate.

The E–P side of core 12 was then propagated to genes using Equation (17). Because the E–P score of a core element is its E–P factor multiplied by a scalar core value, the resulting gene ranking is a property of the E–P factor (here EP03), while the cell factor identifies the cell-type contrast to which that E–P program is coupled. All 16,239 genes obtained nonzero scores: 9,797 positive and 6,442 negative. The degree correction reduced the Spearman correlation between gene degree and absolute score from 0.355 for the uncorrected E–P sum to 0.060 after correction, indicating that the ranking was not driven mainly by the number of candidate enhancer links per gene.

### 2.5 Core-12 Gene Ranking Identifies Contrasting Functional Programs

Of the 16,239 ranked genes, 14,022 (86.35%) mapped to the mouse annotation database used for GO analysis. Preranked GO Biological Process GSEA returned 5,404 terms. At FDR ≤ 0.05, 109 terms had positive NES and 20 had negative NES.

Positive-NES terms were dominated by synaptic function (Table 2), including vesicle-mediated transport in synapse, regulated exocytosis, neurotransmitter secretion, regulation of membrane potential, and modulation of chemical synaptic transmission. Redundancy reduction at Wang semantic similarity 0.70 reduced the 109 significant positive terms to 61 representatives. Across six leading representative processes, recurrent leading-edge genes included *Fmr1*, which appeared in all six selected processes, and *Cacna1a*, *Rims2*, *Rims4*, *Pclo*, *Rimbp2*, *Stx1a*, *Stx1b*, *Grik5*, and *Atp2a2*, each appearing in five of the six. These results define a coherent presynaptic neurotransmission/vesicle-release program.

**Table 2.** Representative GO Biological Process enrichments for core 12.

| **GO Biological Process** | **NES** | **FDR** |
|---|---|---|
| Vesicle-mediated transport in synapse | 2.108 | 1.21 × 10<sup>−6</sup> |
| Regulated exocytosis | 2.050 | 6.73 × 10<sup>−6</sup> |
| Regulation of membrane potential | 1.862 | 6.73 × 10<sup>−6</sup> |
| Neurotransmitter secretion | 2.144 | 6.73 × 10<sup>−6</sup> |
| Transport along microtubule | 2.087 | 1.81 × 10<sup>−5</sup> |
| Synaptic membrane adhesion | −2.472 | 8.72 × 10<sup>−5</sup> |
| Notochord development | −2.171 | 2.13 × 10<sup>−3</sup> |
| Blood vessel morphogenesis | −1.575 | 8.49 × 10<sup>−3</sup> |
| Presynaptic membrane organization | −2.019 | 2.18 × 10<sup>−2</sup> |

The strongest negative term was synaptic membrane adhesion (NES = –2.472, FDR = 8.72×10*^−^*^5^). Other negative representatives included presynaptic membrane organization and multiple developmental or morphogenetic processes. Leading-edge genes shared by at least two of the six leading negative representatives included *Wnt5a* (five processes), *Fgfr2* (four), *Fgf10*, *Gli2*, and *Sox9* (three each), and *Gli3*, *Egfr*, *Ptn*, *Tcf7l2*, *Hgf*, *Jag1*, and *Rbpj*. Because these signaling genes are reused across many developmental GO annotations, terms such as notochord or digestive-tract morphogenesis were interpreted as a general developmental/morphogenetic signaling module rather than literal tissue programs.

The overlap structure of the leading-edge genes is shown in Figure 3. The positive program is dominated by a dense presynaptic module in which many genes recur across most representative GO processes. By contrast, the negative program is more heterogeneous and contains partially overlapping developmental, morphogenetic, and synaptic adhesion gene sets.

**Fig. 3.**
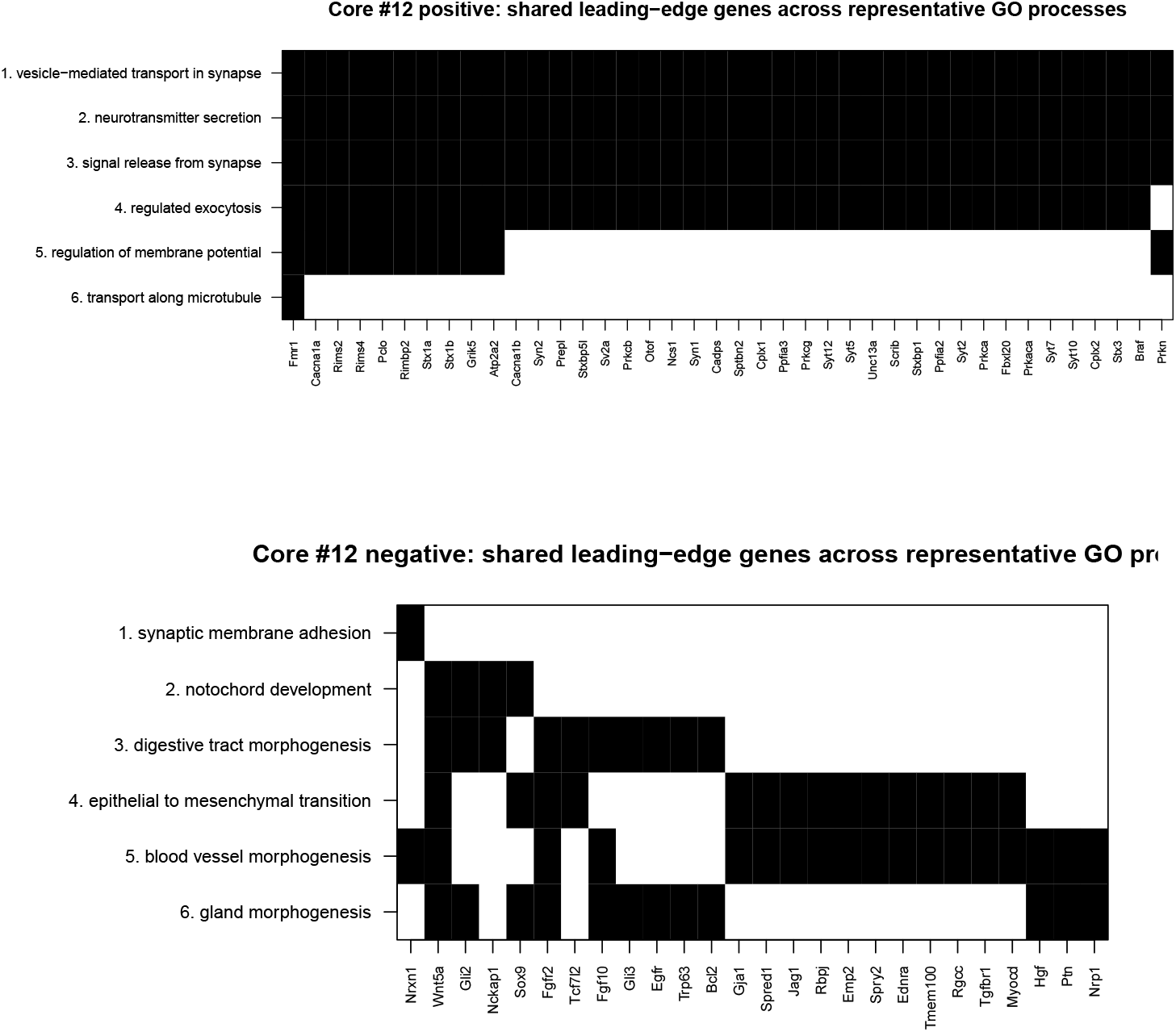
Shared leading-edge genes across representative GO Biological Process terms for core 12. The upper panel shows the positive-NES program and the lower panel the negative-NES program after semantic-similarity-based GO redundancy reduction. Rows are representative GO processes and columns are leading-edge genes. Black cells indicate that a gene belongs to the leading edge of the corresponding process. The positive program contains a densely shared presynaptic neurotransmission/vesicle-release gene module, whereas the negative program shows a more heterogeneous developmental/morphogenetic and synaptic-adhesion structure.

### 2.6 Backprojection Reveals Strong Enhancer-Chromatin and 3D Signals

The latent programs were examined directly in the original measurements. Because the cells (the extremes of CELL03) and the genes (from the core-12 ranking) were selected using a decomposition of these same measurements, the comparisons below are descriptive: they show how the selected axis appears in each original modality, and their *P* values are not tests of independent hypotheses. For each gene of a module, RNA was its own expression, whereas ATAC, H3K27ac, and 3D proximity were taken from the candidate E–P pairs of that gene whose core-12 score had the sign of the module: the ATAC and H3K27ac signals of the paired enhancers and the 3D distances of those E–P pairs, each averaged within the gene before averaging across genes (Methods). For the positive synaptic module, CELL03-positive cells showed markedly higher ATAC and H3K27ac than CELL03-negative cells (Table 3). The rank-biserial effects were 0.921 for ATAC and 0.906 for H3K27ac, with replicate-adjusted FDR values below 10*^−^*^15^. The transformed 3D proximity signal –log(distance) was also higher in CELL03-positive cells. By contrast, the positive-module RNA mean showed only a small, nonsignificant difference at the module level.

**Table 3.** Original-modality backprojection for the positive and negative gene modules.

| **Module** | **Modality** | **Mean CELL03+** | **Mean CELL03−** | **Rank-biserial** |
|---|---|---|---|---|
| Positive | RNA logCP10K | 0.7817 | 0.7308 | 0.140 |
| Positive | ATAC logCP10K | 0.07868 | 0.04770 | 0.9208 |
| Positive | H3K27ac logCP10K | 0.05019 | 0.02256 | 0.9064 |
| Positive | − log(3D distance) | −1.3977 | −1.4608 | 0.4808 |
| Negative | RNA logCP10K | 0.3336 | 0.4006 | −0.6112 |
| Negative | ATAC logCP10K | 0.07589 | 0.06243 | 0.3912 |
| Negative | H3K27ac logCP10K | 0.03548 | 0.02548 | 0.4040 |
| Negative | − log(3D distance) | −1.2778 | −1.4309 | 0.7816 |

The negative developmental/morphogenetic module displayed a different relationship among modalities. Module-level RNA was clearly lower in CELL03-positive cells (rank-biserial –0.611), and ATAC and H3K27ac were moderately higher (0.391 and 0.404), whereas the strongest effect was increased 3D proximity in CELL03-positive cells (0.782). Thus, physical proximity did not map monotonically to transcriptional output.

Figure 4 shows the standardized module scores for the 100 extreme CELL03-loading cells ordered by decreasing CELL03 loading. The chromatin and 3D patterns are visible directly at the cell level: the positive module exhibits higher ATAC and H3K27ac toward the CELL03-positive side, whereas the negative module shows a particularly strong 3D-proximity shift.

**Fig. 4.**
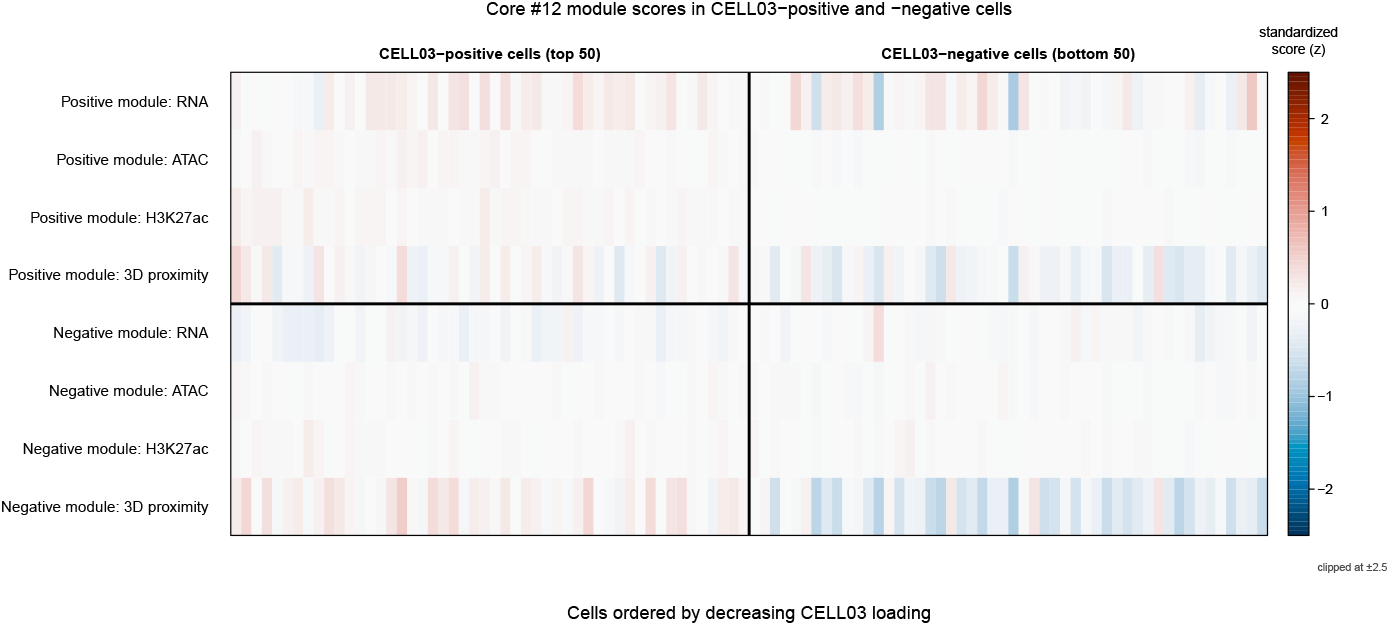
Backprojection of core-12 gene modules to the original four modalities across CELL03 extreme-loading cells. Columns are the 100 cells selected from the two extremes of CELL03 and ordered by decreasing CELL03 loading; the dashed line separates the positive- and negative-loading groups. Rows show standardized RNA, ATAC, H3K27ac, and 3D proximity scores for the positive and negative gene modules, on a diverging scale centred at zero (red, above the mean; blue, below the mean; values in the colour bar). The figure illustrates that the core-12 axis is expressed most strongly in chromatin accessibility, H3K27ac, and 3D organization rather than as a uniform module-wide RNA shift.

Gene-level effects supported these module-level results. Examples included higher positive-cell RNA for *Prkn* and *Syn2* (rank-biserial 0.82 and 0.49); higher positive-cell ATAC and H3K27ac for synaptic genes including *Stx1a*, *Syn2*, *Ncs1*, and *Stxbp1* (0.45– 0.57, FDR≤4×10*^−^*^4^); and strong 3D proximity effects among negative-module genes such as *Wnt5a*, *Rbpj*, *Jag1*, *Gli2*, *Gli3*, and *Hgf* (0.47–0.88, FDR ≤ 1.1×10*^−^*^4^). The 3D observation coverage, defined for each cell as the fraction of a module’s genes for which an E–P 3D distance was observed in that cell, averaged 0.995–1.000 across module-by-group combinations, arguing against differential 3D missingness as the explanation.

### 2.7 Multimodal Contrast Is Reproducible across R1, R2, and R3

The standardized positive-module minus negative-module contrast showed strong differences between CELL03-positive and CELL03-negative cells for ATAC, H3K27ac, and 3D. In the pooled replicate-adjusted analysis, rank-biserial effects were 0.514 for ATAC, 0.552 for H3K27ac, and –0.742 for 3D; RNA showed a smaller effect (0.328). The 100 extreme CELL03 cells were distributed as 21 positive/17 negative in R1, 7/23 in R2, and 22/10 in R3. Despite this imbalance, all four modalities had the same effect direction in all three replicates (Table 4). ATAC and H3K27ac were positive in each replicate, whereas the 3D contrast was negative in each replicate. The signed Stouffer combination yielded FDR values of 2.50×10*^−^*^4^ for ATAC, 1.54×10*^−^*^5^ for H3K27ac, and 1.90×10*^−^*^8^ for 3D. RNA was directionally concordant with a smaller combined effect (FDR = 0.023).

**Table 4.** Replicate-wise rank-biserial effects for the standardized positive-module minus negative-module contrast.

| **Modality** | **R1** | **R2** | **R3** | **Signed-Stouffer FDR** |
|---|---|---|---|---|
| RNA | 0.333 | 0.391 | 0.200 | 0.0233 |
| ATAC | 0.501 | 0.217 | 0.691 | 2.50 × 10<sup>−4</sup> |
| H3K27ac | 0.608 | 0.379 | 0.682 | 1.54 × 10<sup>−5</sup> |
| 3D | −0.776 | −0.913 | −0.564 | 1.90 × 10<sup>−8</sup> |

The replicate-wise effect pattern is summarized visually in Figure 5. The positive-module ATAC and H3K27ac effects remain positive in R1, R2, and R3, while the positive-minus-negative 3D contrast remains negative for all three replicates. Detailed boxplots for the eight module-by-modality combinations and replicate-wise effect trajectories are provided in Supplementary Figures S1 and S2, respectively.

**Fig. 5.**
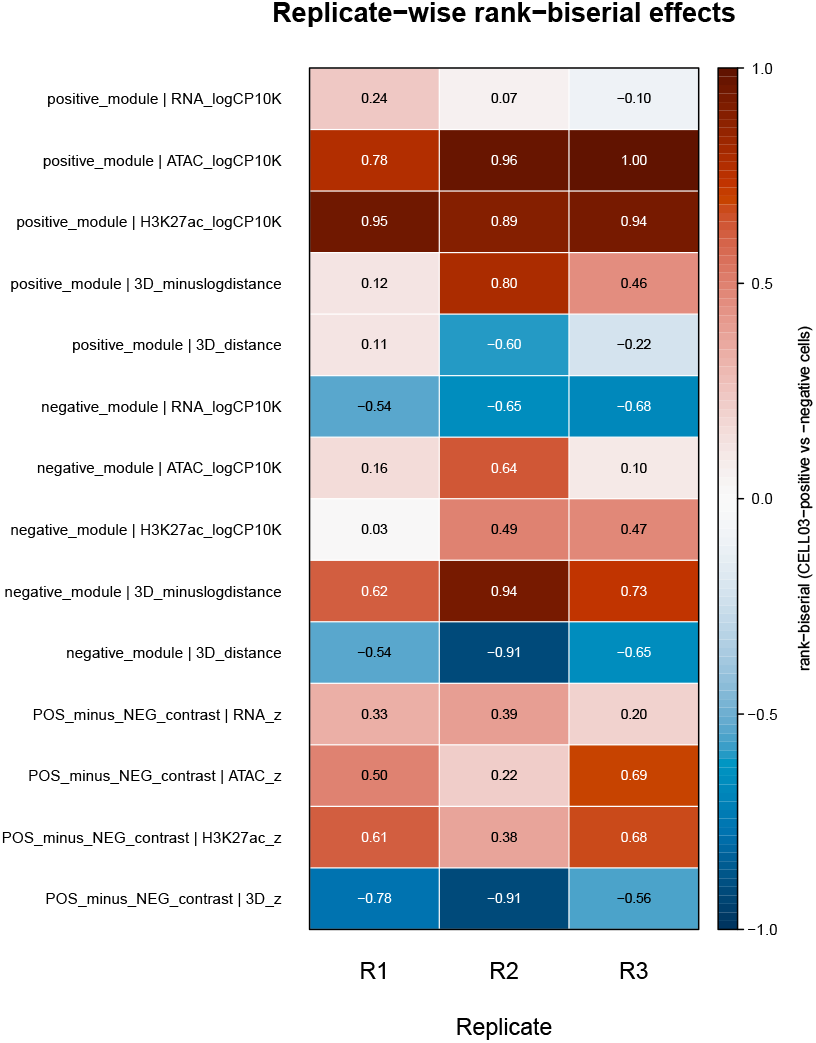
Replicate-wise rank-biserial effects for original-modality backprojection and the positive-minus-negative module contrast. Columns correspond to biological replicates R1–R3. Rows show rank-biserial effect sizes comparing CELL03-positive with CELL03-negative cells for the positive module, negative module, and their standardized contrast; the value is printed in each cell, and the colour bar shows the scale (red, higher in CELL03-positive cells; blue, lower). The consistent direction across replicates is especially evident for positive module ATAC/H3K27ac and for the 3D contrast.

### 2.8 Core 12 Is Recovered by Independent Reanalysis of Each Pair of Replicates

The backprojection above uses factors learned from all three replicates. To test whether core 12 depends on any one replicate, the entire analysis, from stage-1 SVD to stage-2 HOSVD, was repeated independently on each two-replicate subset (R1+R2, R1+R3, R2+R3) with unchanged code, and the factor vectors were compared directly with those of the full data (Supplementary Table S3). The E–P factor of core 12 (EP03) was recovered at the same position in every subset (|*r*| = 0.971–0.973 over all E–P pairs; the next-best subset factor had |*r|* = 0.05–0.06), and so was its cell factor CELL03 (|*r|* = 0.91–0.99 over shared cells). The core element at the position of core 12 in each subset gave gene rankings that agreed with the full data (Spearman *ρ* = 0.974–0.984; 79–92 of the top 100 genes on each side shared) and GO results that agreed as well (Jaccard index of significant positive-NES terms 0.60–0.62; Spearman correlation of NES over 5,404 common terms 0.94–0.96), with the same leading terms in every run: vesicle-mediated transport in synapse on the positive side and synaptic membrane adhesion on the negative side. An automatic rule that picks the largest core element containing the cell factor that best separates Ndnf/Lamp5 from L3/4 IT cells selected core 12 in the full data but a nearly tied neighbouring element in the subsets; the factor comparison shows that this reflects near-ties among core values rather than a change in the factors.

Stage-1 components with nearly equal singular values are defined only up to rotation within their subspace. For every block of stage-1 singular values with a relative gap below 0.05, the singular vectors were rotated within the block (20 random rotations, and a 45*^◦^* rotation for two-component blocks) and stage 2 was recomputed (Supplementary Table S2). The element matching core 12 was found at the same position, (3, 3, 2, 1), after every rotation, and its gene ranking was unchanged (Spearman *ρ ≥* 0.9999).

### 2.9 The Program Requires the Integration of Modalities

To test whether core 12 could be obtained from fewer modalities, stage 2 was recomputed for each single modality and for each leave-one-modality-out combination, and the factors of each variant were matched directly with the E–P and cell factors of core 12 (Supplementary Table S6). No single modality reproduced the program. Cell factors resembling CELL03 existed in single-modality decompositions (for example, |*r|* = 0.94 for ATAC alone), but the core element that coupled them to the best-matching E–P factor was zero (|*G|*≈10*^−^*^17^): in a single-modality decomposition each E–P factor is tied to the cell factor of its own stage-1 component, and the E–P and cell patterns of core 12 do not belong to the same stage-1 component in any modality. Among the leave-one-out variants, removing H3K27ac preserved both the gene ranking (EP03 matched with |*r|* = 0.96; Spearman *ρ* = 0.94) and the GO enrichment (Jaccard index 0.51; NES correlation 0.88), consistent with redundancy between H3K27ac and accessibility. Removing RNA preserved the E–P ranking (|*r|* = 0.98; *ρ* = 0.85) but removed the GO enrichment (7 versus 109 significant positive-NES terms). Removing ATAC or 3D changed both (*ρ* = 0.33 and 0.40). The functionally coherent program of core 12 thus depends jointly on accessibility, chromatin contact, and expression.

### 2.10 Prioritized E–P Pairs Agree with the Cell-Type-Resolved Links of the Original CHARM Study

The candidate E–P pairs are deliberately broad. To assess whether the pairs prioritized by core 12 agree with enhancer–gene links inferred by other means, they were compared with the 4,954 cell-type-resolved links reported in the original CHARM study [3], which were obtained from the same data by an independent method (Poisson regression with Shapley attribution) and were not used in the decomposition (Supplementary Figure S4, Supplementary Table S5). Of these links, 906 (18%) overlapped a candidate enhancer of the same gene in our candidate set. Links that CHARM assigned only to Ndnf/Lamp5 cells and those assigned only to L3/4 IT cells received core-12 scores of opposite sign (73% versus 2.5% positive; rank-biserial *r*_rb_ = 0.89, Wilcoxon *P* = 1.6×10*^−^*^19^), in the orientation of the CELL03 contrast. Accordingly, Ndnf/Lamp5 links were enriched among the 1,000 highest-scoring positive-side pairs (odds ratio 37; Cochran–Mantel–Haenszel test stratified by enhancer–TSS distance decile, *P* = 4.3×10*^−^*^59^) and absent from the top negative-side pairs, whereas L3/4 IT links showed the converse pattern (odds ratio 179, *P* = 1.2×10*^−^*^258^; none on the positive side). The same directions held when the comparison was restricted to genes with at least one CHARM link (odds ratios 8.8 and 39). Ranked by absolute score, CHARM links of any cell type were enriched beyond what distance explains (odds ratio 24), as were activity-by-contact-like links computed from the pseudobulk data (odds ratio 20.1). Distance alone, however, ranked both link sets better than core 12 did, as expected for a component that represents a cell-type contrast rather than a link predictor. Because both reference sets derive from the same data set, these results show concordance between analyses rather than independent validation.

### 2.11 Recurring E–P Factors among the Largest Core Elements

Nineteen E–P factors occurred among the 100 largest core elements; EP01 and EP03 were the most frequent (10 elements each; Supplementary Table S7). Their gene rankings defined different programs: EP01 separated synapse-organization genes (284 significant negative-NES terms), EP03 carried the presynaptic program of core 12, and EP07 separated synaptic genes from myelination and immune genes. They also differed in the genomic distance of their top-scoring pairs: EP01 favoured distal pairs (median 534 kb versus 468 kb for all pairs, *P* = 1.1×10*^−^*^10^), EP07 proximal pairs (410 kb, *P* = 2.0×10*^−^*^7^), and EP03 showed no preference (*P* = 0.95). Some E–P factors were shared across several cell-type contrasts: EP03 combined with four cell factors, contrasting astrocytes and oligodendrocytes with neurons as well as inhibitory interneurons with L3/4 IT neurons. Others occurred with a single cell factor only, for example EP16 with a microglia-versus-olfactory-bulb factor.

## 3 Methods

### 3.1 Data Source and Genome Annotation

Processed and reconstructed data from the CHARM study were obtained from NCBI GEO accession GSE303006 [3]. The analysis used mouse brain total RNA counts, ATAC fragment files, H3K27ac fragment files, reconstructed 20-kb 3D structures, and cell-level CHARM metadata. According to the GEO record, cell identifiers beginning with R1, R2, and R3 correspond to the three mouse brain biological replicates. The source study used GRCm38/mm10 and GENCODE mouse release M23; the same GENCODE M23 annotation was used here.

### 3.2 ATAC and H3K27ac Peak Calling and Enhancer Definition

ATAC and H3K27ac fragment files were converted to BED format and pooled across brain cells for peak calling with MACS3 (version 3.1.0; -g mm). ATAC peaks were called using --nomodel --shift -50 --extsize 100 --keep-dup all -q 0.01. H3K27ac peaks were called in broad-peak mode using --nomodel --shift -50 --extsize 100 --keep-dup all --broad --broad-cutoff 0.05.

Promoters were defined based on GENCODE M23 gene transcription start sites as TSS ±1 kb. Candidate active enhancers were defined as ATAC peaks overlapping a H3K27ac broad peak after removing peaks overlapping the promoter intervals. Candidate enhancer–gene pairs were generated by linking each enhancer center to all annotated gene TSSs within 1 Mb on the same chromosome.

RNA genes were required to have a nonzero count in at least 1% of the original RNA cells. Gene symbols that mapped ambiguously to more than one GENCODE gene identifier were excluded from the E–P candidate set.

### 3.3 Enhancer, 3D, and Cell Quality Control

Enhancer-level detection fractions were calculated across the common ATAC/H3K27ac/3D cell set. Enhancers were retained when both ATAC and H3K27ac were detected in at least 0.5% of cells. This yielded 42,669 enhancers after the E–P and 3D filters.

Enhancer centers and promoter TSSs were assigned to 20-kb genomic bins. For each cell, 3D Euclidean distance between the enhancer and promoter bins was calculated from the reconstructed CHARM 3D coordinates. Maternal and paternal distances were calculated separately when available and averaged for the unphased analysis. A 20-kb bin pair was retained when 3D distance was available in at least 90% of the common cells (pair-level 3D coverage ≥0.90) and the distance variance was finite. This pair-level coverage is distinct from the cell-level 3D observation coverage used in the backprojection. After 3D quality control, 529,900 of 555,189 candidate bin pairs were usable.

At the cellular level, cells were required to have at least 250 ATAC enhancer-associated fragments and at least 100 H3K27ac enhancer-associated fragments. For RNA, cells were required to have a library size of at least 500 counts and 500 detected genes. After intersection with cells having the required 3D data, 4,258 cells were retained for the final four-modality analysis.

### 3.4 Normalization

For ATAC and H3K27ac, cell depth was calculated using all enhancer candidate rows, and the retained enhancer matrix was transformed as

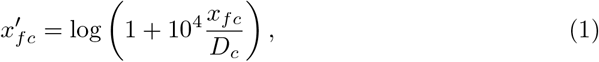

where *x_fc_* is the fragment count for feature *f* in cell *c* and *D_c_* is the corresponding total depth. The RNA matrix was normalized to the same log-CP10K scale. Each retained feature was then centered and scaled across cells.

For 3D, positive finite Euclidean distances *d_bc_* were converted to proximity-like values

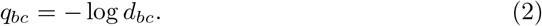

For each 20-kb bin pair, *q_bc_* was standardized over observed cells. Missing standardized 3D values were set to zero, so that missingness corresponded to the feature mean after standardization.

### 3.5 Stage-1 Modality-Specific Decomposition

Each modality was reduced independently to *K* = 20 components. The rank *K* = 20 was set as a single round value for all modalities before the analysis; it was not derived from a formal criterion and was not tuned to the results. It is a parameter of the model: a different *K* defines a different decomposition. The variance retained by this truncation is reported for every modality (Supplementary Table S1). For ATAC, H3K27ac, and RNA, truncated SVD was applied to the feature-wise standardized matrix. The decomposition was expressed as

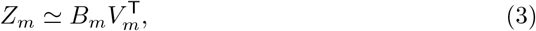

where *B_m_* contains modality-specific feature scores and *V_m_* contains orthonormal cell loadings. ATAC and H3K27ac feature rows correspond to enhancers, RNA rows to genes, and 3D rows to 20-kb E–P bin pairs.

For the 3D matrix, a blockwise randomized SVD was used with oversampling 10, one power iteration, and random seed 20260811. For deterministic orientation, the sign of each component was chosen such that the cell loading with the largest absolute magnitude was positive. The retained rank-20 reconstruction of each modality was subsequently scaled to Frobenius norm one. Consequently, no modality dominated stage 2 solely because of a larger overall numerical scale.

The *k*th component was interpreted only as the *k*th singular-value-ranked component *within* a modality. No Procrustes rotation was used to force one-to-one correspondence between components from different modalities. A sign-only orientation check was performed before stage 2. The index *k* therefore serves as a bookkeeping index and does not assume that the *k*th components of different assays describe the same process. Because the E–P, cell, and modality factors of stage 2 are shared across all *k*, a process carried by component 2 of one assay and component 5 of another is represented by the same pair of E–P and cell factors, with core values in both slices. Aligning components across assays would require a matching criterion chosen before the decomposition; the rank index avoids that choice and leaves the cross-modality combination to stage 2.

### 3.6 E–P Mapping and Degree Correction

Let *p*denote the final E–P pairs, and let *e*(*p*), *g*(*p*), and *b*(*p*) denote the enhancer, target gene, and 20-kb bin pair assigned to E–P pair *p*. Because one enhancer, gene, or 20-kb bin pair can appear in multiple E–P rows, their degrees were defined as

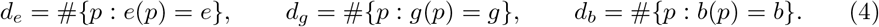

Inverse square root weights 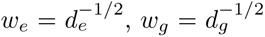, and 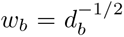 were used. These weights satisfy, for example,

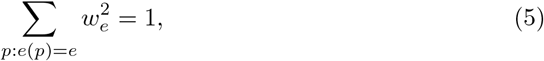

so replicating an underlying feature over several E–P rows does not inflate its total squared contribution.

### 3.7 Implicit Stage-2 Tensor

The conceptual stage-2 tensor was

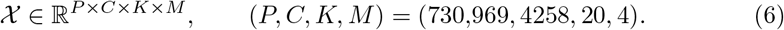

Its entries were defined as follows:

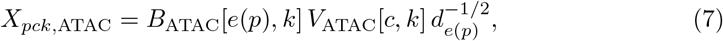

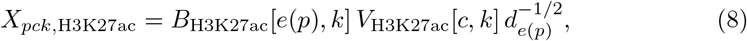

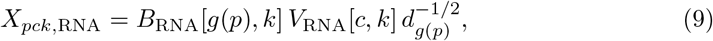

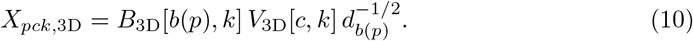

The full tensor was never materialized.

### 3.8 Implicit HOSVD

HOSVD [6] was performed using Tucker ranks 20 20×20×4 for the E–P, cell, stage-1-component, and modality modes, respectively.

For the E–P mode, the 80 pair-side vectors (20 components *×* 4 modalities) were conceptually concatenated into a *P ×* 80 matrix *A*. Instead of forming the *P ×P* Gram matrix, the exact 80 *×* 80 Gram matrix

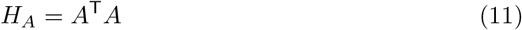

was accumulated in blocks. If *H_A_* = *W* Λ*W* ^T^, the retained E—P factor was obtained blockwise as

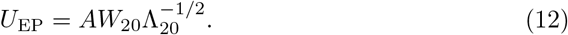

The cell-mode factor was computed from a 4,258×80 matrix whose columns were the modality-specific cell loadings weighted by the norm of the corresponding pair-side vector. The original stage-1 *K* = 20 ordering was deliberately preserved, so the component-mode factor was the identity rather than a further rotation of *K*. The modality factor was obtained from the 4 *×* 4 modality-mode Gram matrix.

The core tensor *G*, of size 20 *×* 20 *×* 20 *×* 4, was computed directly from projected pair and cell vectors and the modality factor. Its element *G_abkr_* links the *a*th E–P factor (the *a*th column of *U*_EP_, denoted EP*a*), the *b*th cell factor (CELL*b*), the *k*th stage-1 component (the component factor being the identity, *k* is the same index as in Equation (10)), and the *r*th modality factor (MOD*r*). Core elements are ranked by |*G_abkr_*|. The retained multilinear energy fraction was defined as the squared Frobenius norm of the core divided by the squared Frobenius norm of the conceptual tensor. The explicit formula for every factor and for the core, including the degree weights, is derived in Supplementary Note 1.

Two properties of the construction are proved there. First, because the degree weights satisfy Equation (5), each pair-side column has the norm of the corresponding stage-1 feature-score column, and the cell factors span the same subspace as the leading left singular vectors of the concatenated, norm-weighted stage-1 cell loadings, that is, of a PCA of the vertically concatenated stage-1 reconstructions. The E–P pairing therefore enters the result through the E–P factors and the core tensor, not through the cell subspace. Second, the decomposition is invariant to the ordering of E–P pairs, cells, and modalities. Because each factor matrix of the HOSVD is obtained from its own Gram matrix, the leading factors and the core values among them do not change when a larger stage-2 rank is retained.

### 3.9 Cell-Type and Replicate Association of Cell Components

The CHARM metadata were matched to the final 4,258 cells. Replicate labels R1–R3 were obtained from the cell identifiers, consistent with the GEO metadata [3]. For each distinct cell HOSVD component appearing among the 100 largest absolute Tucker-core elements, the component loading was compared across annotated cell types and replicates using the Kruskal–Wallis test.

Effect size was summarized as

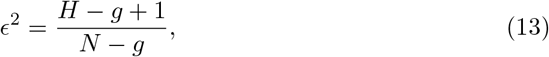

where *H* is the Kruskal–Wallis statistic, *g* is the number of groups, and *N* is the number of cells. For interpretability, the 50 cells with the largest positive and 50 with the largest negative loading for each component were tested for cell-type enrichment by one-sided Fisher exact tests.

To assess whether cell-type effects generally exceeded replicate effects, one point was generated for each distinct cell component represented among the 100 largest core entries, comparing 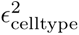 with 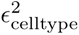. A one-sided sign test against probability 0.5 is reported in the Supplementary Information as a descriptive summary only, because the 12 components are derived from the same data and their association statistics are not independent.

Because cell-type composition differs between replicates, cell type and replicate were also assessed jointly. The cell-type-by-replicate composition was tabulated and tested by a *χ*^2^ test. For each of the 12 components, the loading was modelled by the linear model

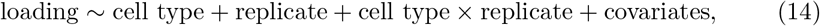

where the covariates were log ATAC fragment count, log H3K27ac fragment count, log RNA library size, log number of detected genes, and the fraction of 3D bin pairs observed in the cell. For each term, the partial effect size was

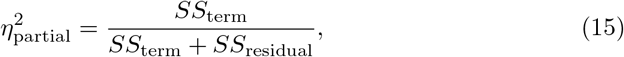

where *SS*_residual_ is the residual sum of squares of the full model and *SS*_term_ is the type-II sum of squares of the term: cell type and replicate were each adjusted for the other main effect and the covariates, the interaction was adjusted for both main effects and the covariates, and each covariate was adjusted for all other terms. The interaction term therefore measures how much replicate effects differ between cell types beyond what the two main effects explain. The model was fitted to raw and to rank-transformed loadings. Replicate effects within each cell type were tested by Kruskal–Wallis tests for cell types with at least 20 cells in at least two replicates, with Benjamini–Hochberg adjustment over all tests. For CELL03, the Ndnf/Lamp5 versus L3/4 IT contrast was computed within each replicate (rank-biserial *r*_rb_), and the largest difference between replicates within a cell type was expressed relative to the difference between the two cell types.

### 3.10 Tracing Core 12 to E–P Pairs and Genes

Core elements were ranked by absolute value. Core rank 12, (*a, b, k, r*) = (3, 3, 2, 1) was selected as a representative example because its cell component, CELL03, exhibited a strong cell-type association and weak replicate association.

For every E–P pair *p*, the signed E–P-side score associated with this core element was

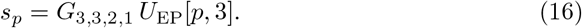

To produce a signed ranking of all 16,239 genes, a degree-corrected target-gene score was then calculated from all 730,969 E–P pairs as follows:

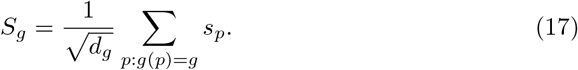

Because *s_p_* is the E–P factor column multiplied by the scalar *G_abkr_*, the gene ranking derived from a core element depends only on its E–P factor *a*, up to the sign of *G_abkr_*; the cell factor *b* identifies the cell-type contrast with which that E–P program is coupled. In the full data, CELL03 is higher in Ndnf/Lamp5 than in L3/4 IT cells and the modality factor MOD01 has a positive sum, so positive scores correspond to the Ndnf/Lamp5 side. In the reanalyses described below, every run was oriented by the same rule (cell factor higher in Ndnf/Lamp5 cells; modality factor with a positive sum, or, for variants with a different set of modalities, a positive inner product with the core-12 modality factor over the shared modalities).

### 3.11 GO Biological Process GSEA and Redundancy Reduction

The degree-corrected gene ranking was analyzed by preranked mouse Gene Ontology (GO) Biological Process GSEA using clusterProfiler::gseGO [7]. Gene symbols were mapped with org.Mm.eg.db; gene sets containing 10–500 genes were included, and a Benjamini–Hochberg false discovery rate (FDR) ≤ 0.05 was used for significance.

Positive- and negative-NES terms were processed separately. Redundancy was reduced with GO semantic similarity using GOSemSim [8] and the Wang graph-based similarity measure [9]. Significant GO terms were ordered by FDR and then by absolute NES; a term was assigned to an existing representative group if its semantic similarity to the representative was at least 0.70. Otherwise, it initiated a new group.

For biological module construction, the six highest-ranking representative GO processes on each side were considered. Leading-edge genes appearing in at least two of those six processes were retained (up to 40 genes if necessary) to define positive and negative modules.

### 3.12 BackProjection to the Original Four Modalities

CELL03-positive and CELL03-negative cells were defined as the 50 largest and 50 smallest CELL03 loadings, respectively. For the positive gene module, only E–P pairs with a positive core-12 E–P score were used; for the negative gene module, only E–P pairs with a negative core-12 E–P score were used.

RNA was evaluated at the gene level. For ATAC and H3K27ac, all sign-consistent enhancers linked to each module gene were first averaged within that gene; gene-level values were then averaged with equal gene weight. The same procedure was used for 3D bin pairs. This within-gene then across-gene averaging prevented genes with many candidate E–P links from dominating the module score.

Group differences between CELL03-positive and CELL03-negative cells were summarized using the Wilcoxon rank-sum test and rank-biserial effect size. Because both the cells and the gene modules were selected from a decomposition of the same data, these statistics describe the selected axis and are not independent tests. Replicate-adjusted effects were estimated using the linear model.

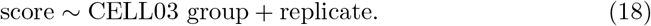

### 3.13 Replicate-Wise Reproducibility

The 100 CELL03 extreme-loading cells were split into R1, R2, and R3. For each replicate, the positive-module minus negative-module contrast was calculated for standardized RNA, ATAC, H3K27ac, and 3D scores. Direction, rank-biserial effect size, and Wilcoxon *P* values were recorded separately.

For the descriptive combined test, two-sided Wilcoxon *P* values were converted to signed normal scores using the sign of the replicate-specific rank-biserial effect. Signed Stouffer statistics were combined with weights 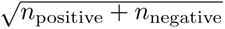 and corrected by the Benjamini–Hochberg procedure across all 14 module-by-modality and contrast summaries. Because the HOSVD factors were learned using all three replicates, this analysis was interpreted as within-dataset reproducibility rather than independent validation.

### 3.14 Reanalysis of Two-Replicate Subsets

For each two-replicate subset, the stage-1 and stage-2 scripts were rerun unchanged on the cells of those replicates, with the same E–P pairs. Factors were compared with those of the full data by |Pearson *r*| : E–P factors over all 730,969 E–P pairs and cell factors over the shared cells; for each full-data factor, the best- and second-best-matching subset factors were recorded. The core element at the position of core 12, (*a, b, k, r*) = (3, 3, 2, 1) in each run’s own factor order, was analyzed in every run as in the full data (gene ranking and GO-BP GSEA), and agreement was summarized by the Spearman correlation of gene scores, the overlap of the top 100 genes on each side, the Jaccard index of significant GO terms, and the correlation of NES over common terms.

### 3.15 Rotations within Near-Degenerate Stage-1 Subspaces

Within each modality, consecutive stage-1 singular values with a relative gap below 0.05 were grouped into blocks. Writing the stage-1 factors as *B_m_* = *U_m_* diag(*d̄_m_*) and *V_m_*, the left and right singular vectors of each block were rotated by the same orthogonal matrix, with the singular values unchanged, so that the reconstruction changed only through the small differences of the singular values within the block. All blocks of all modalities were rotated simultaneously, by 20 random orthogonal matrices drawn from the Haar distribution and, for two-component blocks, also by a 45*^◦^* rotation, and stage 2 was recomputed each time. The core element matching core 12 (by maximal |*r|* of the E–P and cell factors) and the agreement of its gene ranking with the original were recorded. Exact permutations of the component index leave the stage-2 E–P and cell factors unchanged (Supplementary Note 1).

### 3.16 Modality Ablation

Stage 2 was recomputed, with the same stage-1 results, for each single modality and each leave-one-modality-out combination. Because the E–P factors are linear combinations of the stage-1 pair-side columns (*U*_EP_ = *AW* Λ*^−^*^1*/*2^), their correlations with the four-modality E–P factors were accumulated blockwise over E–P pairs without storing the factors. In each variant, the E–P and cell factors best matching EP03 and CELL03 were identified, the core element combining them with the largest |*G*| over components and modality factors was taken, and its gene ranking and GO-BP GSEA were compared with core 12.

### 3.17 Comparison with Supported Enhancer–Gene Links

Two reference sets were used. First, the cell-type-resolved enhancer–gene links of the original CHARM study (Supplementary Table 7 of [3]; 4,954 unique pairs at 5-kb resolution with presence in 16 cell types) were matched to candidate pairs when a candidate enhancer overlapped a listed 5-kb bin for the same gene symbol. Second, activity-by-contact-like scores [10] were computed from pseudobulk data: activity as the geometric mean of mean ATAC and H3K27ac log-CP10K over cells, contact as the mean over cells of the inverse 3D distance of the bin pair, and the score as their product normalized over the candidate enhancers of each gene; pairs with a score of at least 0.02 were considered supported. For each reference set, the 1,000 and 5,000 highest-scoring pairs on the positive side, the negative side, and by absolute value were compared with the remaining pairs by Cochran–Mantel–Haenszel tests stratified by deciles of enhancer–TSS distance, and the average precision of each ranking was compared with 200 permutations of the scores within distance deciles and with the average precision of distance alone. For the CHARM links, the analysis was repeated within genes having at least one CHARM link, and the core-12 scores of links specific to Ndnf/Lamp5 and specific to L3/4 IT cells were compared by the Wilcoxon rank-sum test.

### 3.18 Characterization of Cell and E–P Factors among the Largest Core Elements

For each cell factor among the 100 largest core elements, the loadings were standardized over cells and averaged within each annotated cell type. For each E–P factor, the number of top-100 core elements containing it and the recurring combinations of E–P and cell factors were counted. For the most frequent E–P factors, the gene ranking (Equation 17) and GO-BP GSEA were computed from the largest core element containing the factor, and the enhancer–TSS distances of the 1,000 pairs with the largest absolute score were compared with all pairs by the Wilcoxon rank-sum test.

### 3.19 Use of AI Tools

Large language models were used as assistants in this study. For the original version of the manuscript, GPT-astra was used to help design the analyses, to write and test the analysis code, and to draft the manuscript text; the stage-1 rank *K* = 20 was adopted from a value proposed during this code development and was not tuned to the results. For the revision, Claude (Anthropic) was used in the same way: to help design the additional analyses requested by the reviewers, to write and test the corresponding R and Python scripts and batch-job scripts, and to draft the revised text and the response to the reviewers. All analyses were run by the authors on the Kyoto University ICR supercomputer, every numerical result reported in the manuscript was taken from the logs of these runs, which are deposited with the code, and the authors checked all code, results, and text. The AI tools are not authors, and the authors take full responsibility for the content of the manuscript.

## 4 Discussion

This study introduces an implicit hierarchical HOSVD strategy for integrating four single-cell regulatory modalities while preserving their natural feature identities. The principal computational advantage is that the 730,969×4,258×20×4 stage-2 tensor never needs to be stored. Instead, the E–P mode is recovered from an exact 80×80 Gram matrix, and the cell mode from a compact 4,258×80 representation. This makes a multiway E–P analysis feasible even though direct tensor materialization would require an impractically large memory footprint.

The analysis also differs conceptually from integrations that primarily seek a common cell embedding. The stage-1 component axis and modality axis remain explicit, and enhancer, gene, and 3D-bin features are mapped to E–P pairs only after modality-specific decomposition. On the cell side alone, the method is close to conventional integration: because the degree weights preserve the norm of each stage-1 component, the cell factors span the same subspace as a PCA of the concatenated, norm-weighted stage-1 reconstructions (Supplementary Note 1). What the method adds is the core tensor, which couples each cell factor to E–P factors through the E–P pairing. For the same reason, we did not compare the cell factors with those of concatenated PCA or of multi-view factor models such as MOFA+: such a comparison would concern only the cell side, on which the methods agree by construction, whereas these methods provide no E–P factors and no core through which a cell factor is linked to E–P pairs. The modality ablation showed that this coupling is not available from any single modality: the cell factor of core 12 has close counterparts in single-modality decompositions, but the core element linking it to the matching E–P factor vanishes, because each E–P factor of a single-modality decomposition is tied to the cell factor of its own stage-1 component. The program reported here depended jointly on accessibility, chromatin contact, and expression; H3K27ac was largely redundant with accessibility, and RNA, although dispensable for the E–P ranking, was required for its functional coherence. The degree correction further prevents a gene or enhancer from acquiring excessive weight simply because it participates in many candidate E–P pairs. The reduction of the degree–score Spearman correlation from 0.355 to 0.060 in the Core-12 gene ranking provides an empirical check that this correction achieved its intended purpose.

A central concern in unsupervised decomposition is whether an apparently biological component is instead driven by batch or replicate. Here, the cell-type association was not limited to the detailed CELL03 example. Although cell-type composition differed between replicates, a joint model showed that cell type explained more variance than replicate or their interaction for all 12 distinct cell components represented among the 100 largest core elements. This generalization is important because core 12 was selected after establishing that the cell-type-dominant pattern was widespread among the large core elements. Repeating the whole analysis independently on each pair of replicates recovered the E–P and cell factors of core 12 at the same positions, with nearly identical gene rankings and GO results, so the program does not depend on any single replicate.

Core 12 provided a biologically interpretable example of the framework. Its cell axis contrasted Inh_Ndnf/Lamp5 with Ex_L3/4_IT cells, while its E–P axis separated a presynaptic neurotransmission/vesicle-release module from a developmental/morphogenetic and synaptic-adhesion module. The positive module was driven by recurrent genes encoding proteins closely connected to presynaptic release and excitability, including *Cacna1a*, *Rims2*, *Rims4*, *Pclo*, *Rimbp2*, *Stx1a*, and *Stx1b*. The negative module contained Wnt/FGF/Hedgehog-related developmental signaling genes as well as cell-adhesion genes such as *Nrxn1*. Because the gene ranking of a core element is determined by its E–P factor alone, this E–P program (EP03) is not specific to the two cell types: EP03 also appeared with three other cell factors among the largest core elements, contrasting astrocytes and oligodendrocytes with neurons. Core 12 is therefore best read as the coupling of a shared E–P program to the inhibitory-versus-L3/4 IT contrast. Consistently, the enhancer–gene links that the original CHARM study assigned specifically to Ndnf/Lamp5 or to L3/4 IT cells fell on opposite sides of the core-12 E–P ranking.

The original-modality backprojection showed that the latent axis should not be reduced to a conventional RNA-expression signature. For the presynaptic module, the clearest effects were observed for enhancer accessibility and H3K27ac, whereas its module-level RNA difference was small. In particular, the negative developmental/morphogenetic module could show stronger 3D proximity in CELL03-positive cells even when its RNA output was lower. This partial decoupling is biologically plausible because physical enhancer–promoter contact is not equivalent to transcriptional activation. Enhancer activity reflects multiple regulatory dimensions, including chromatin accessibility, histone state, transcription-factor occupancy, and contact frequency [10]. A four-omics analysis can thus reveal combinations that would be hidden in a transcriptome-only analysis.

The replicate analysis strengthened the within-dataset interpretation. ATAC, H3K27ac, 3D, and RNA effects had identical directions across R1, R2, and R3, and the 3D contrast was large in every replicate (rank-biserial 0.56 to 0.91). R2 contained only seven CELL03-positive extreme cells, reducing power for the chromatin assays, but its effect directions remained concordant with R1 and R3.

This study had several limitations. First, all analyses use one data set. The replicate-wise backprojection is not an independent validation because the factors were learned from all three replicates. The two-replicate reanalysis addresses robustness to the replicate structure, but it is not a test in independent data. We did not use a leave-one-replicate-out design in which a held-out replicate is projected onto factors learned from the other two, because such a projection mixes the decomposition with batch differences between experiments; independent decompositions of each replicate pair, compared with one another and with the full data, avoid this. Second, the detailed functional analysis focused on core 12. All cell factors and recurring E– P factors among the 100 largest core elements are now characterized, but the other programs were not analyzed at the same depth. Third, the candidate E–P search was intentionally broad (enhancer–TSS distance ≤ 1 Mb). Therefore, the findings do not indicate that every retained pair is a functional physical interaction. The tensor analysis should be interpreted as prioritizing candidate E–P-associated programs within this candidate space rather than as a definitive E–P loop caller. The prioritized pairs agreed with the cell-type-resolved links of the original CHARM study [3] and with activity-by-contact-like links beyond what genomic distance explains, but both reference sets were derived from the same data, distance alone ranked them better than core 12, and the CHARM links are themselves predictions; this is concordance, not experimental validation. Fourth, the stage-1 rank *K* = 20 is a parameter of the model that was fixed in advance; a different *K* defines a different decomposition, whose components need not coincide with those reported here.

## 5 Conclusions

An implicit hierarchical HOSVD can integrate single-cell ATAC, H3K27ac, RNA, and 3D genome measurements at the level of candidate enhancer–promoter pairs without constructing the full multiway tensor. Applied to CHARM mouse brain data, the dominant cell-mode structure was consistently more strongly associated with cell type than with biological replicate. A representative core linked a neuronal subtype axis to opposing synaptic and developmental/morphogenetic candidate E–P-associated programs; it was recovered in independent analyses of each pair of replicates, could not be obtained from any single modality, and agreed in direction with the cell-type-resolved links of the original CHARM study. Backprojection to the original measurements showed strong and replicate-consistent effects in enhancer accessibility, H3K27ac, and 3D genome organization, whereas RNA differed clearly only for the developmental module. In this data set, the decomposition therefore identifies candidate programs that span several molecular layers, and it provides an interpretable route from a large multi-omic latent representation back to cells, enhancers, genes, and E–P relationships. Whether such programs generalize to other tissues and data sets remains to be tested.

## Supplementary information

The following supporting information accompanies this manuscript: Figure S1, GSE303006_M23_core12_CELL03_original4modality_module_scores_boxplots.pdf, boxplots for positive and negative core-12 modules across RNA, ATAC, H3K27ac, and 3D; Figure S2, GSE303006_M23_core12_CELL03_replicate_reproducibility_rank_biserial_by_replicate. replicate-wise rank-biserial trajectories for the positive module, negative module, and positive-minus-negative contrast; Figure S3, GSE303006_M23_rev26_celltype-by_component_heatmap.pdf, mean standardized loading of each annotated cell type on the 12 cell factors among the 100 largest core elements (11 cells without a cell-type label in the CHARM metadata are shown as a separate row); Figure S4, GSE303006_M23_rev23c_CHARM_concordance.pdf, core-12 scores of candidate pairs overlapping the cell-type-resolved CHARM links and the enrichment of Ndnf/Lamp5 and L3/4 IT links among the top-scoring pairs. Supplementary Note 1 gives the full derivation of the implicit decomposition. Supplementary Tables S1–S7 (Supplementary Tables S1-S7.xlsx, one sheet per table) are the output tables of the analyses added in revision, each labelled with the script that produced it: S1, stage-1 and stage-2 variance retention and reconstruction errors; S2, rotations within near-degenerate stage-1 subspaces; S3, reanalysis of two-replicate subsets; S4, cell-type-by-replicate composition, joint models, and the descriptive sign test; S5, comparison with CHARM and activity-by-contact-like links; S6, modality ablation; S7, recurring E–P factors among the 100 largest core elements. The complete R analysis scripts are available in the code repository; running them on GSE303006 produces all intermediate and output tables, including the top-100 Tucker core annotation, GO/GSEA output tables, and original-modality backprojection tables, together with a log of every run, and tools/build supplementary tables.py assembles Supplementary Tables S1–S7 from these outputs.

## Acknowledgements

Acknowledgements are not compulsory. Where included they should be brief. Grant or contribution numbers may be acknowledged.

Please refer to Journal-level guidance for any specific requirements.

## Declarations

## Ethics approval and consent to participate

Not applicable.

## Consent for publication

Not applicable

## Data availability

The single-cell four-omics data analyzed in this study are publicly available from NCBI GEO under accession GSE303006, with raw data under BioProject PRJNA1284811.

## Materials availability

Not applicable. This study is a secondary computational analysis of publicly available data.

## Code availability

The source CHARM preprocessing code is available from the repository reported by the original study. The R scripts used for the hierarchical tensor analysis in the present work, including all analyses added in revision and job scripts for batch execution, are available at https://github.com/tagtag/TDCHARM; the version corresponding to the first preprint is tagged v1-preprint and the revised version v2-revision.

## Author Contributions

Y.H.T. planned the study and performed the analyses. Y.H.T. and T.T. evaluated the results and wrote and reviewed the manuscript. Conceptualization, data curation, and analysis were performed by Y.H.T. All the authors have read and agreed to the published version of the manuscript.

