## Supplementary material for "Implicit Hierarchical Tensor Decomposition of Single-Cell Four-Omics Data Reveals Cell-Type-Linked Candidate Enhancer–Promoter-Associated Programs": GSE303006_M23_core12_CELL03_original4modality_module_scores_boxplots.pdf

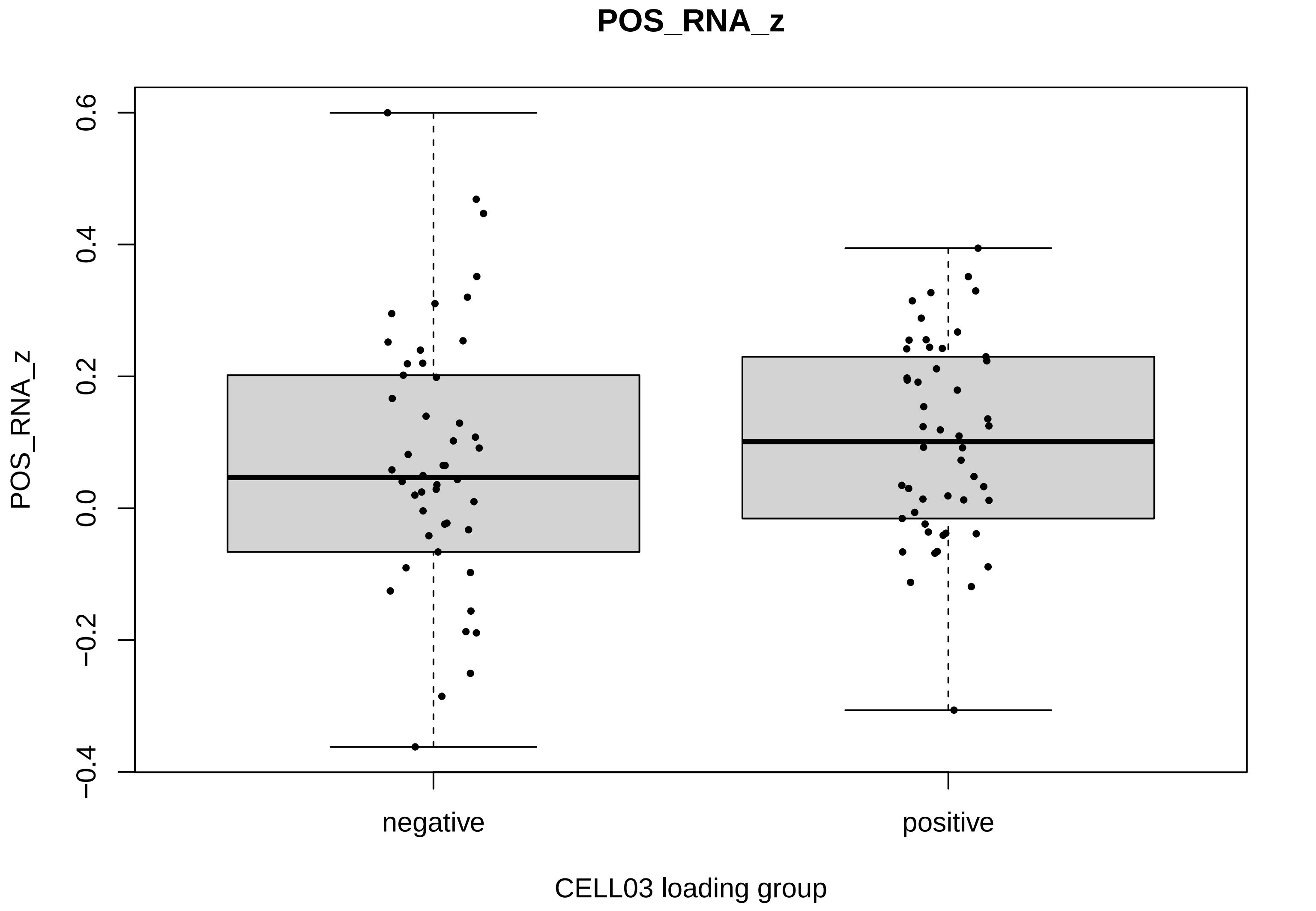
POS_RNA_z
0.6
0.4
0.2
0.0
−0.2
−0.4
POS_RNA_z
negative
positive
CELL03 loading group

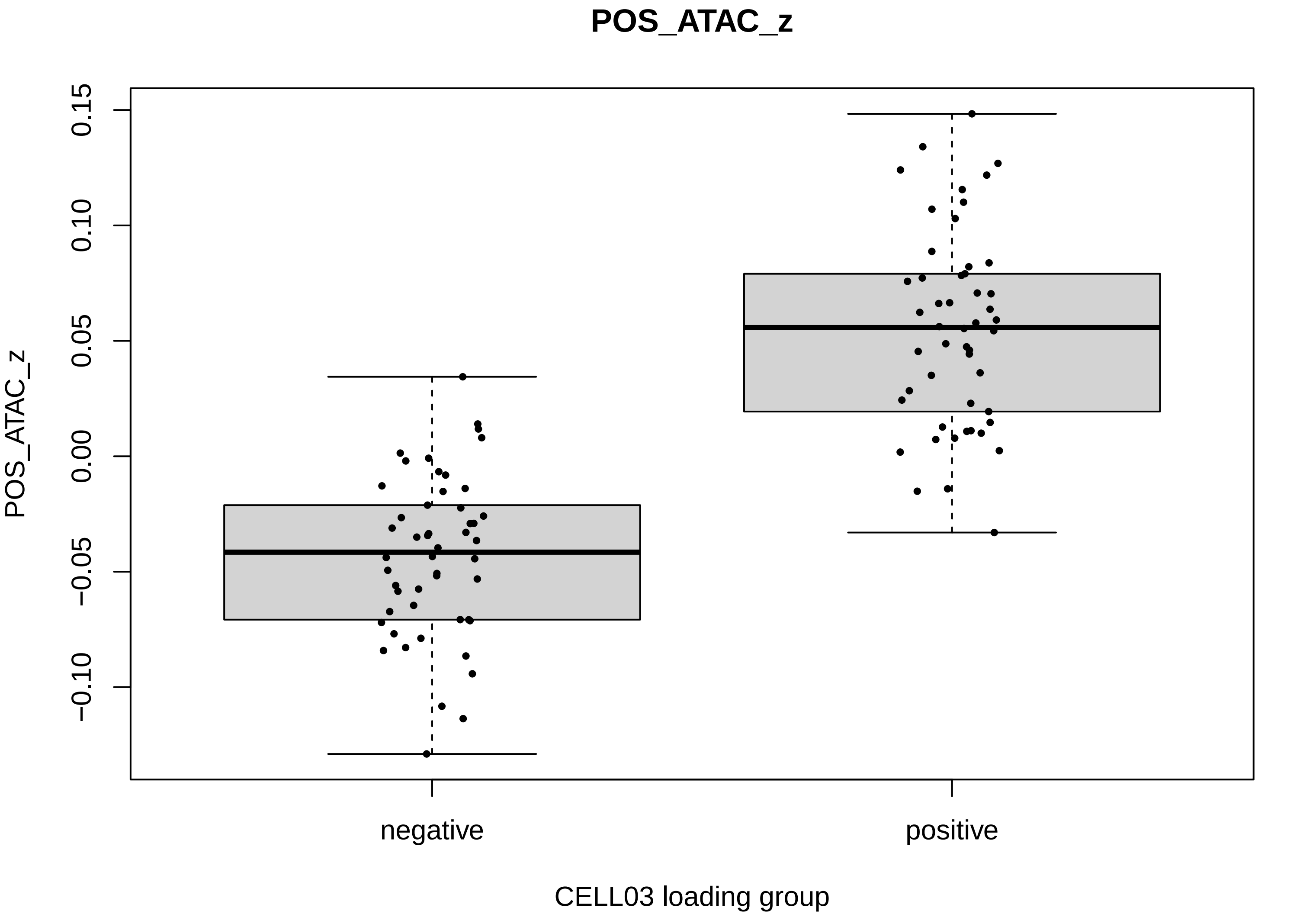
POS_ATAC_z
POS_ATAC_z
0.15
0.10
0.05
0.00
−0.05
−0.10
negative
positive
CELL03 loading group

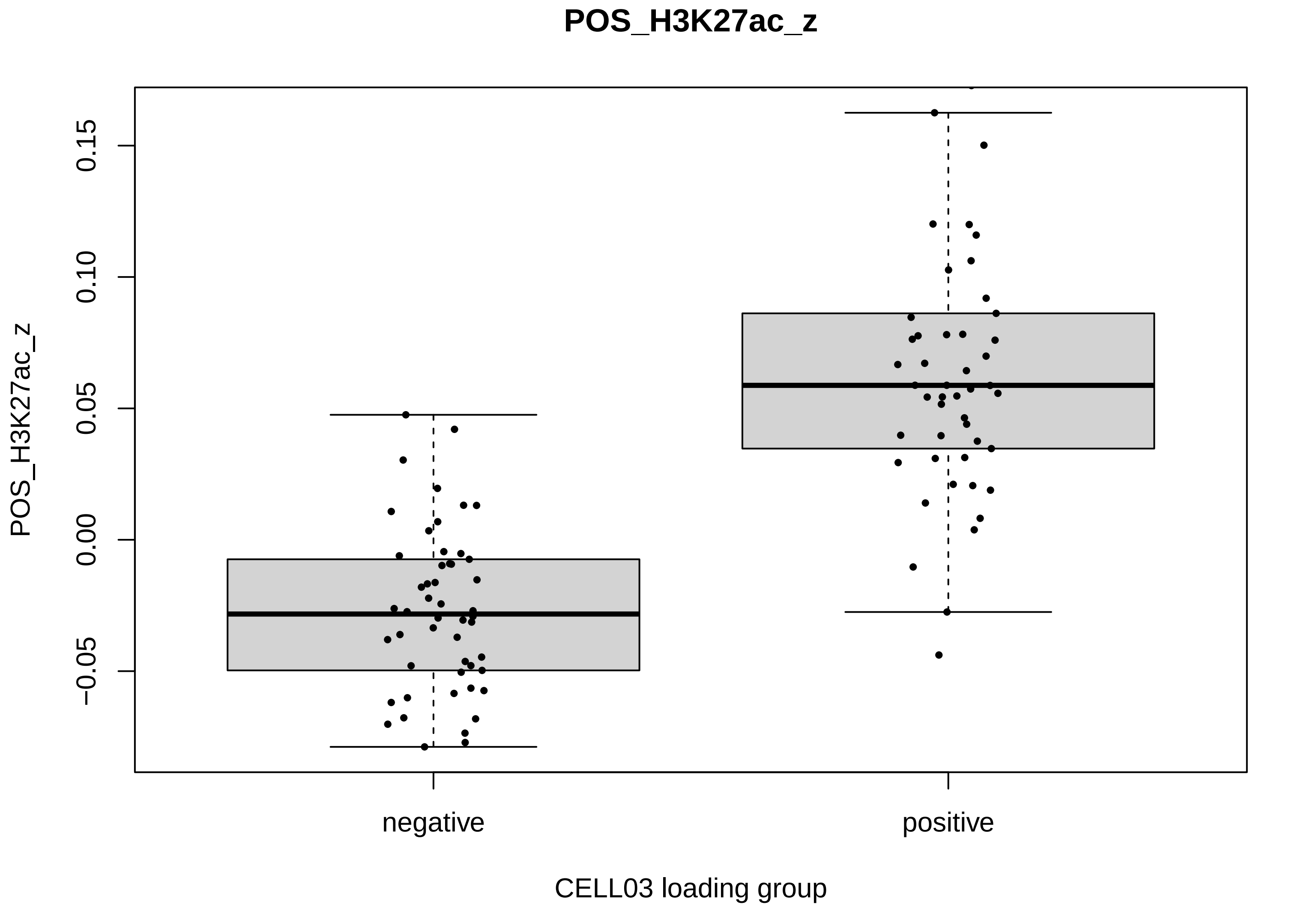
POS_H3K27ac_z
0.15
0.10
0.05
0.00
−0.05
POS_H3K27ac_z
negative
positive
CELL03 loading group

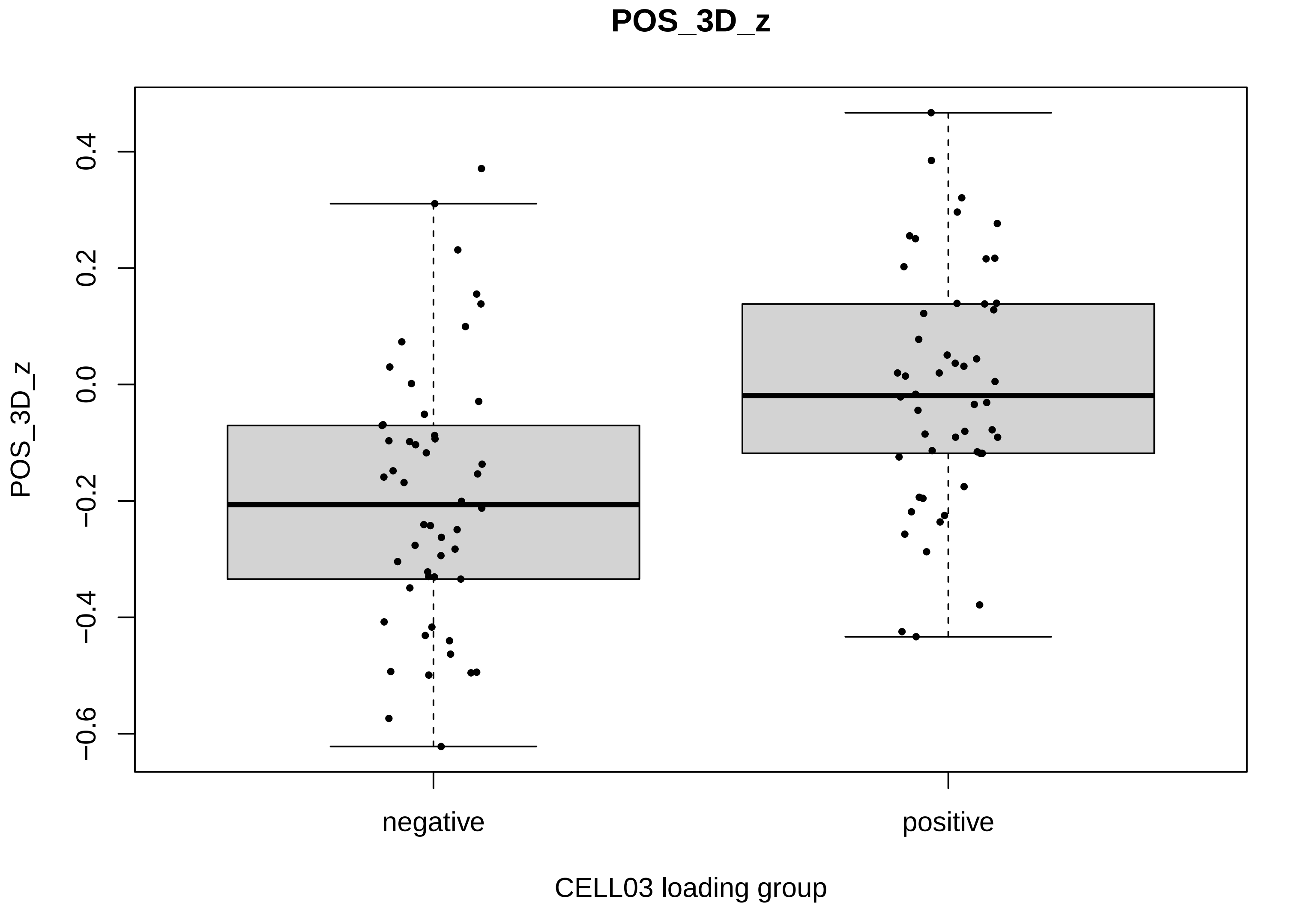
POS_3D_z
POS_3D_z
0.4
0.2
0.0
−0.2
−0.4
−0.6
negative
positive
CELL03 loading group

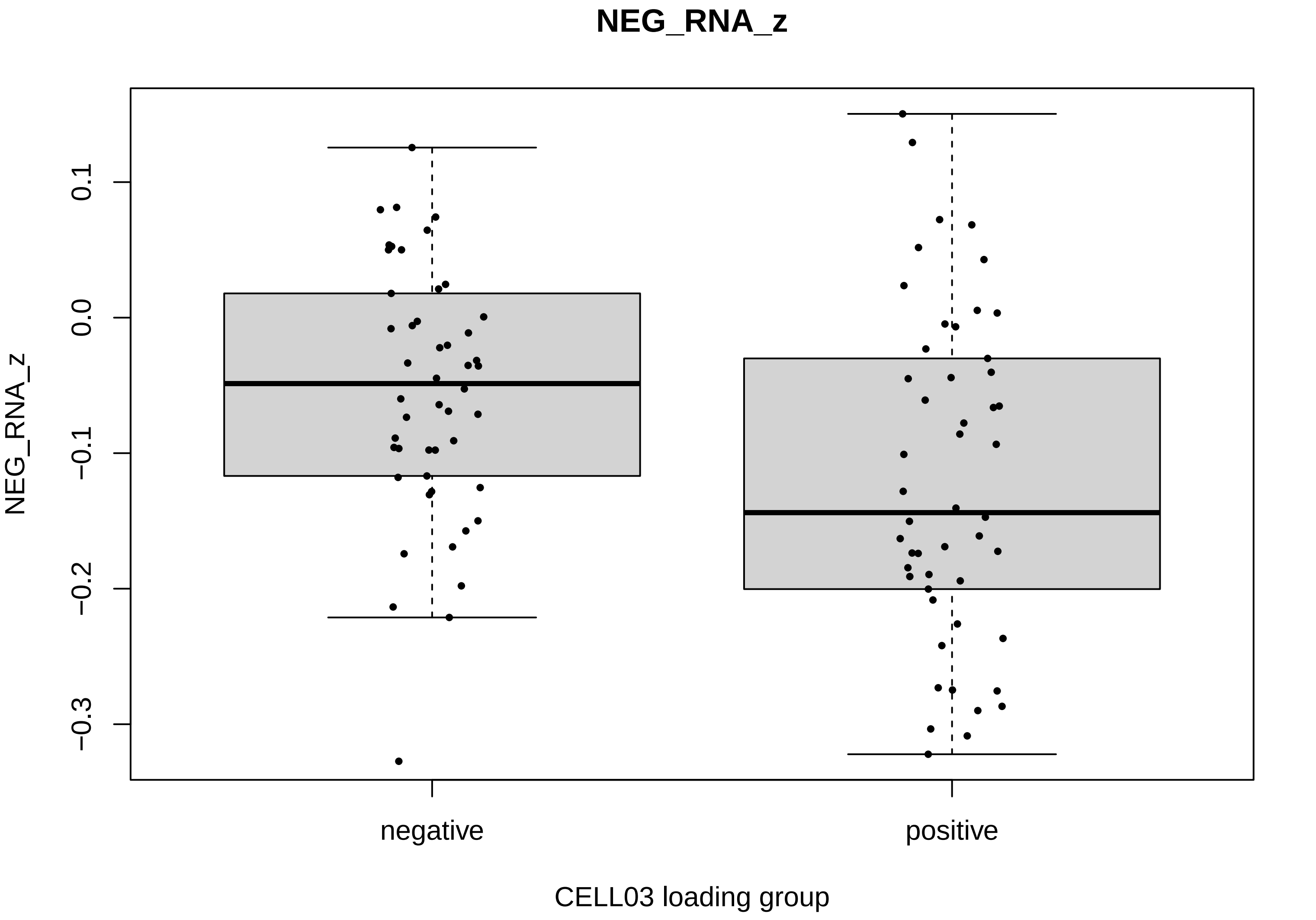

NEG_RNA_z
NEG_RNA_z
0.1
0.0
−0.1
−0.2
−0.3
negative
positive
CELL03 loading group

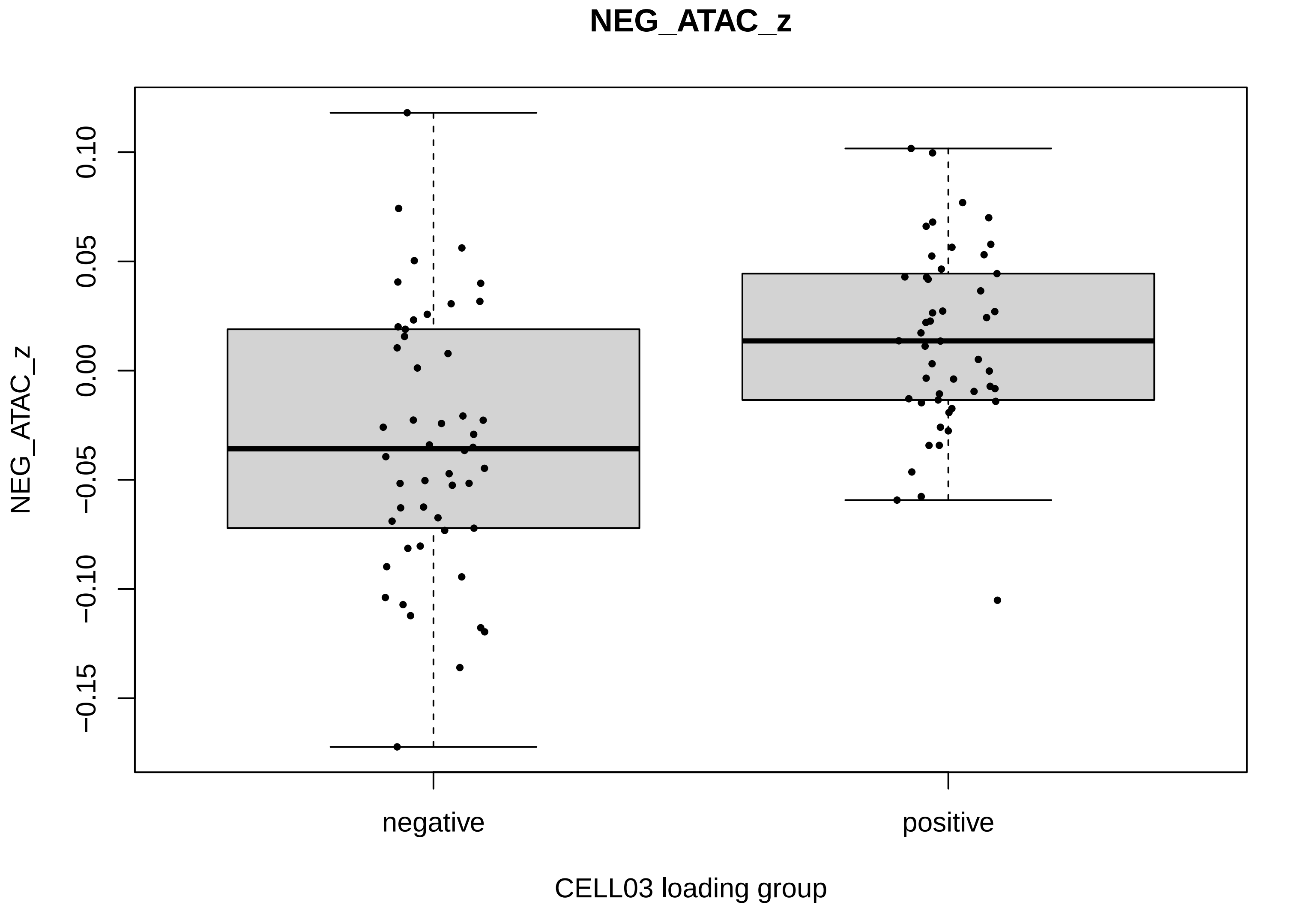
NEG_ATAC_z
NEG_ATAC_z
0.10
0.05
0.00
−0.05
−0.10
−0.15
negative
positive
CELL03 loading group

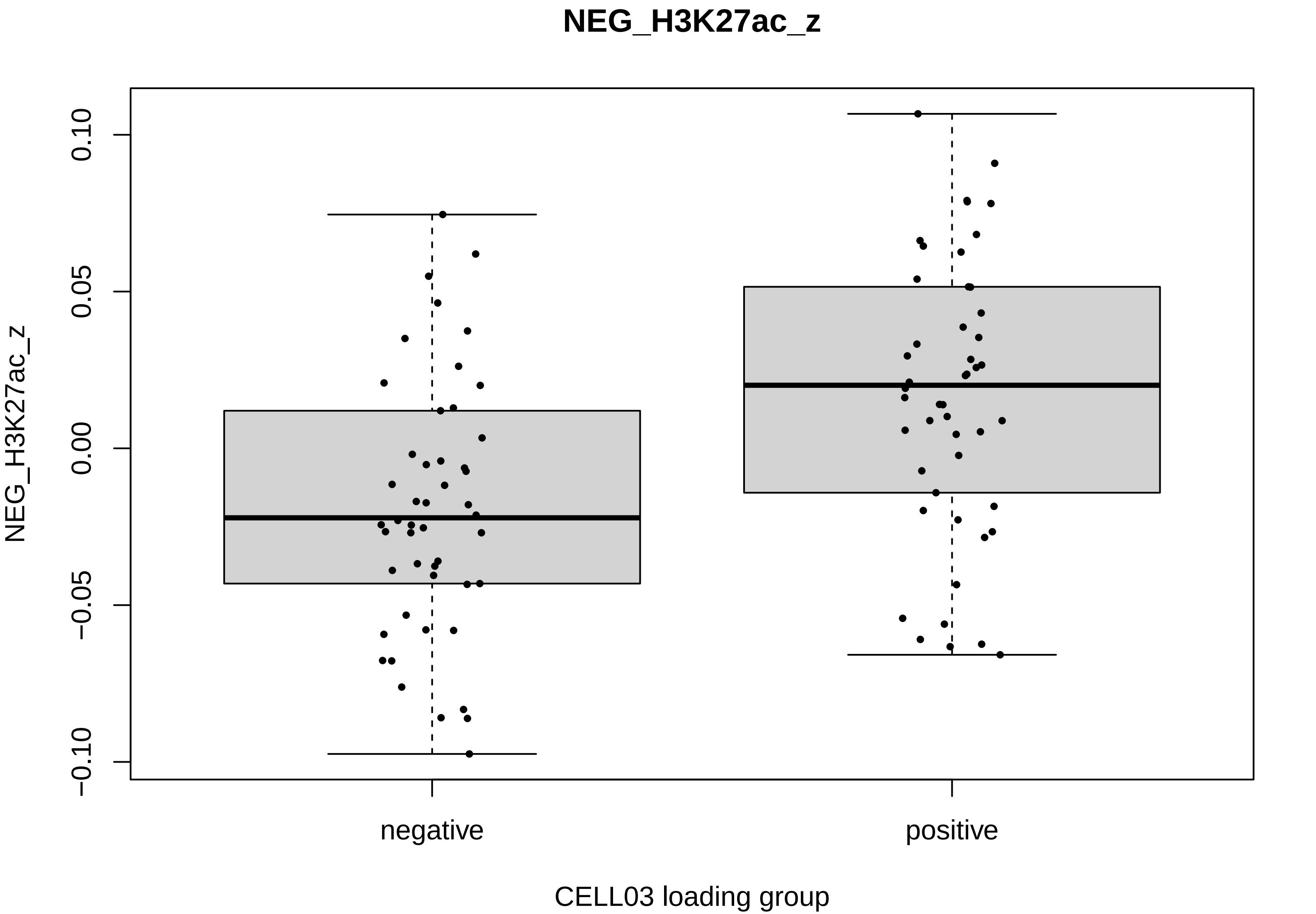
NEG_H3K27ac_z
0.10
0.05
0.00
−0.05
−0.10
NEG_H3K27ac_z
negative
positive
CELL03 loading group

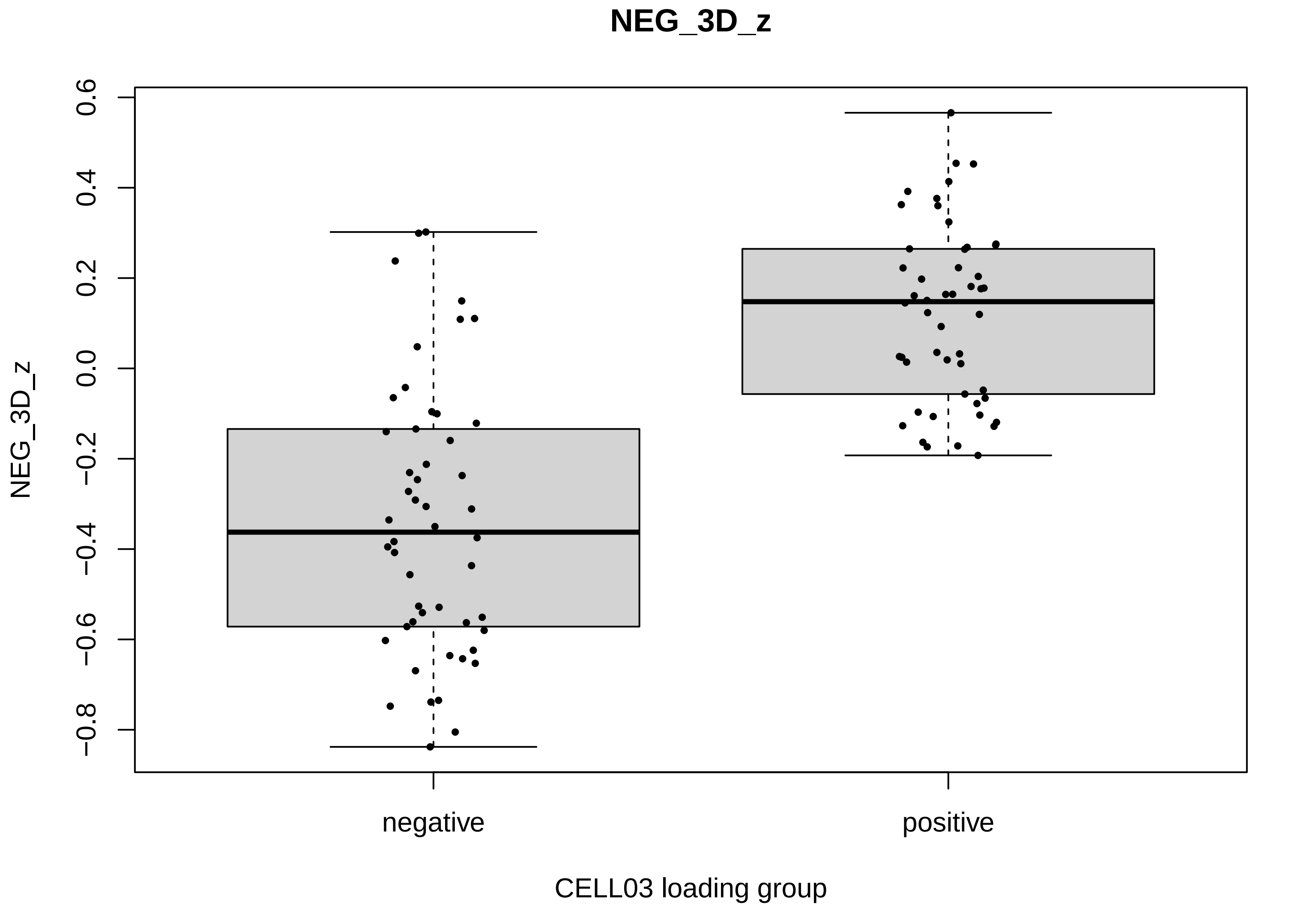
NEG_3D_z
0.6
0.4
0.2
0.0
−0.2
−0.4
−0.6
−0.8
NEG_3D_z
negative
positive
CELL03 loading group
