## Supplementary material for "Implicit Hierarchical Tensor Decomposition of Single-Cell Four-Omics Data Reveals Cell-Type-Linked Candidate Enhancer–Promoter-Associated Programs": GSE303006_M23_core12_CELL03_replicate_reproducibility_rank_biserial_by_replicate.pdf

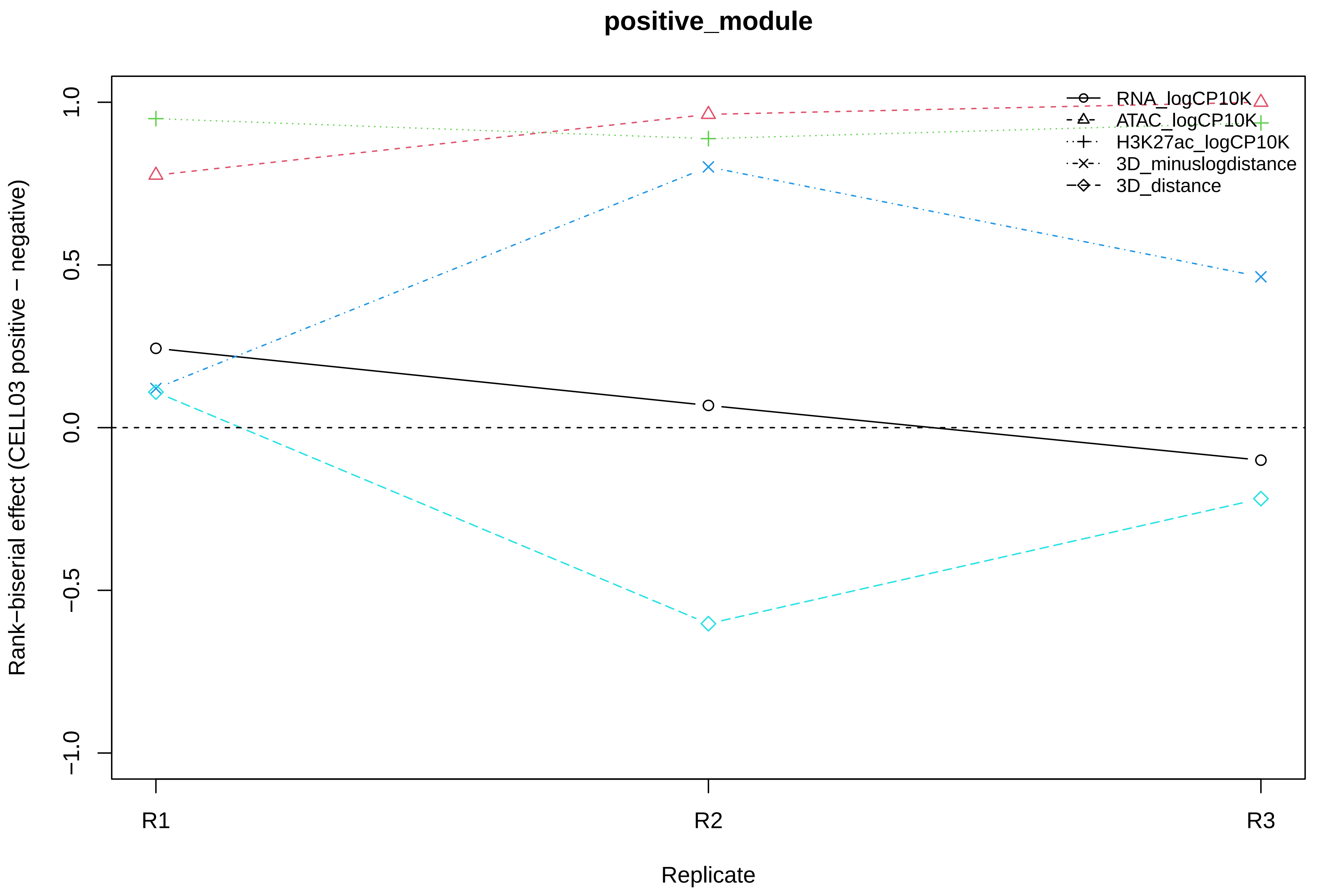

positive_module
Rank−biserial effect (CELL03 positive − negative)
1.0
0.5
0.0
−0.5
−1.0
R1
R2
R3
Replicate
RNA_logCP10K
ATAC_logCP10K
H3K27ac_logCP10K
3D_minuslogdistance
3D_distance

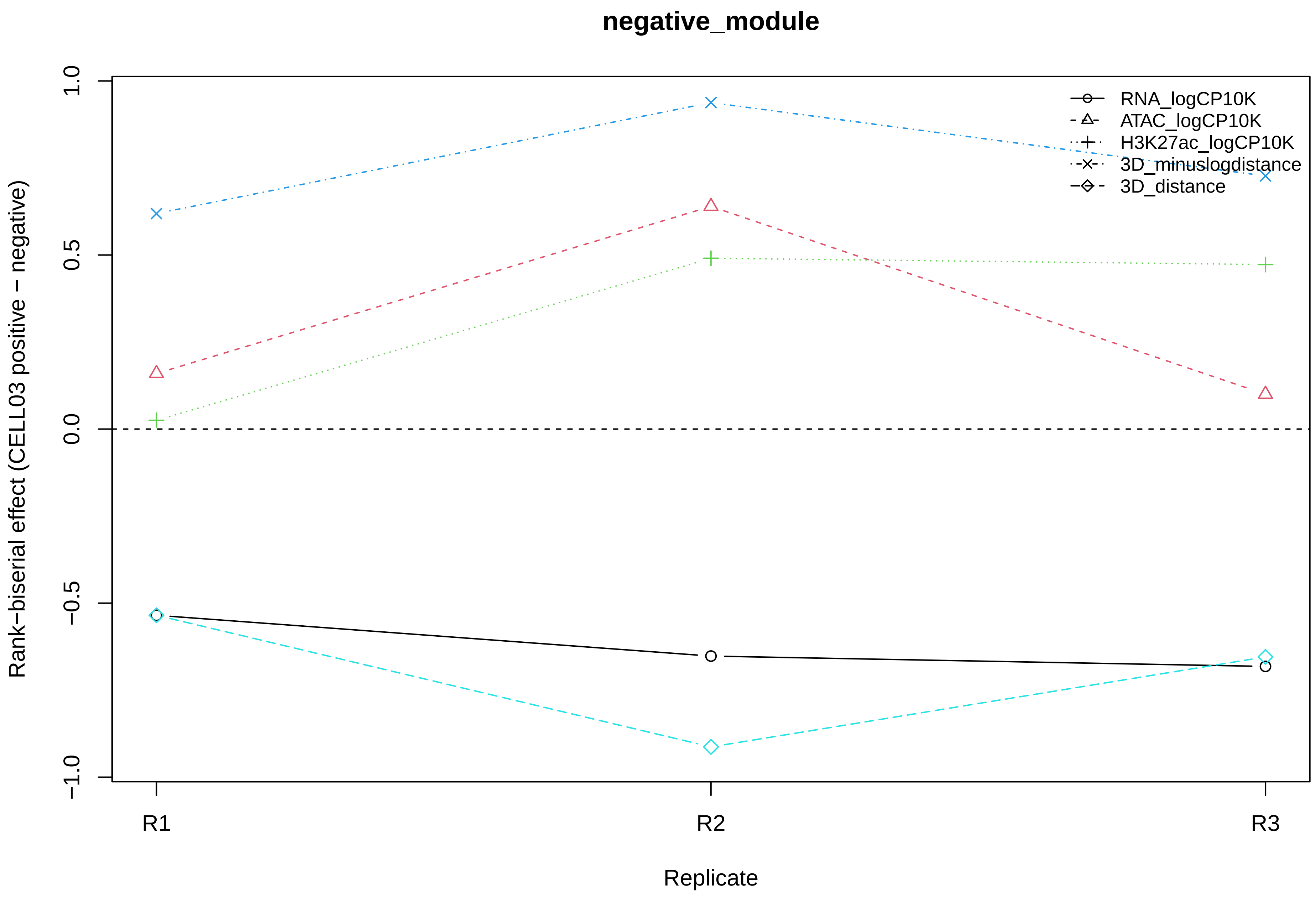
negative_module
Rank−biserial effect (CELL03 positive − negative)
1.0
0.5
0.0
−0.5
−1.0
R1
R2
R3
Replicate
RNA_logCP10K
ATAC_logCP10K
H3K27ac_logCP10K
3D_minuslogdistance
3D_distance

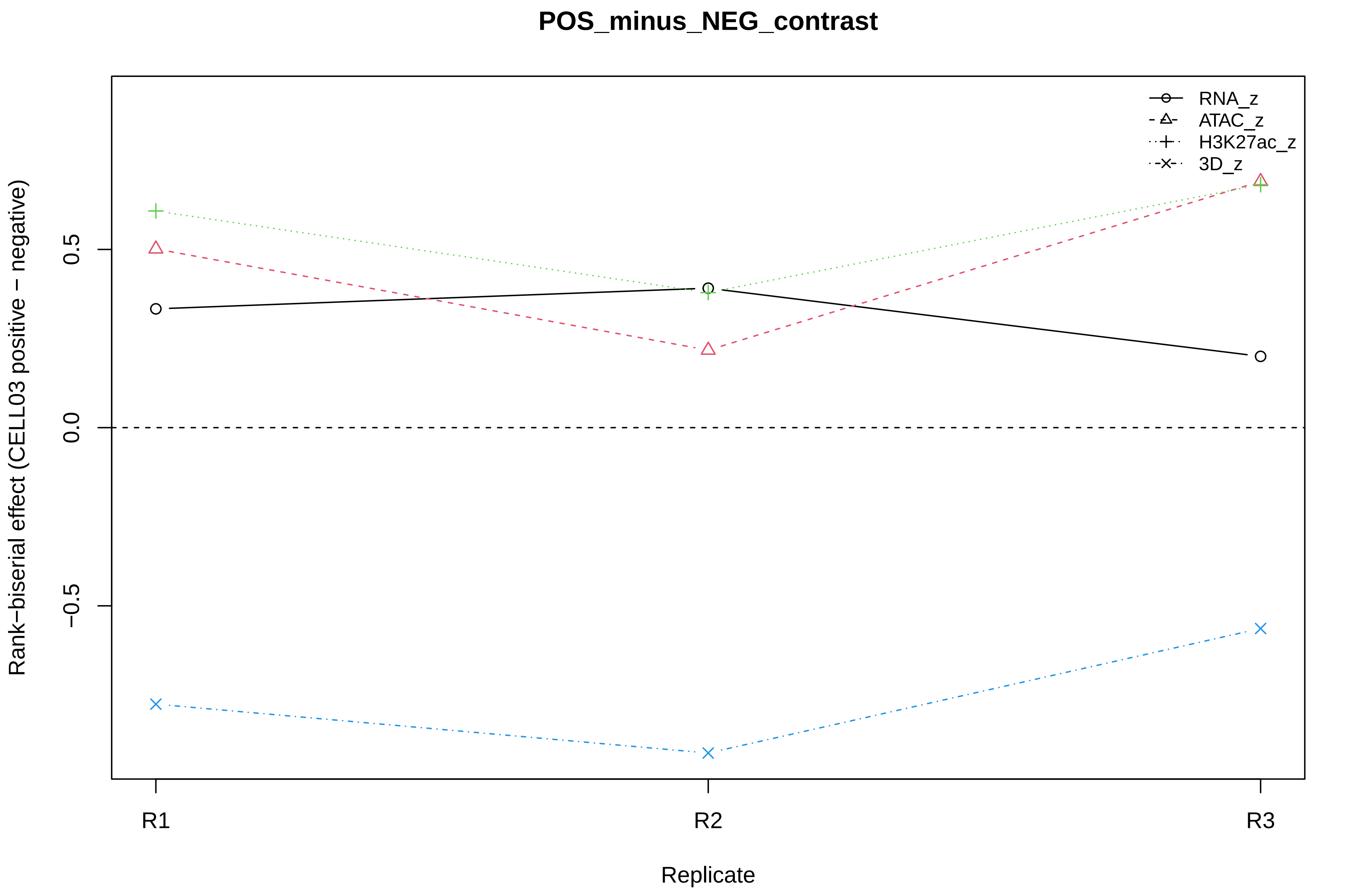
POS_minus_NEG_contrast
RNA_z
ATAC_z
H3K27ac_z
3D_z
0.5
0.0
−0.5
Rank−biserial effect (CELL03 positive − negative)
R1
R2
R3
Replicate
