## Supplementary material for "Implicit Hierarchical Tensor Decomposition of Single-Cell Four-Omics Data Reveals Cell-Type-Linked Candidate Enhancer–Promoter-Associated Programs": GSE303006_M23_rev26_celltype_by_component_heatmap.pdf

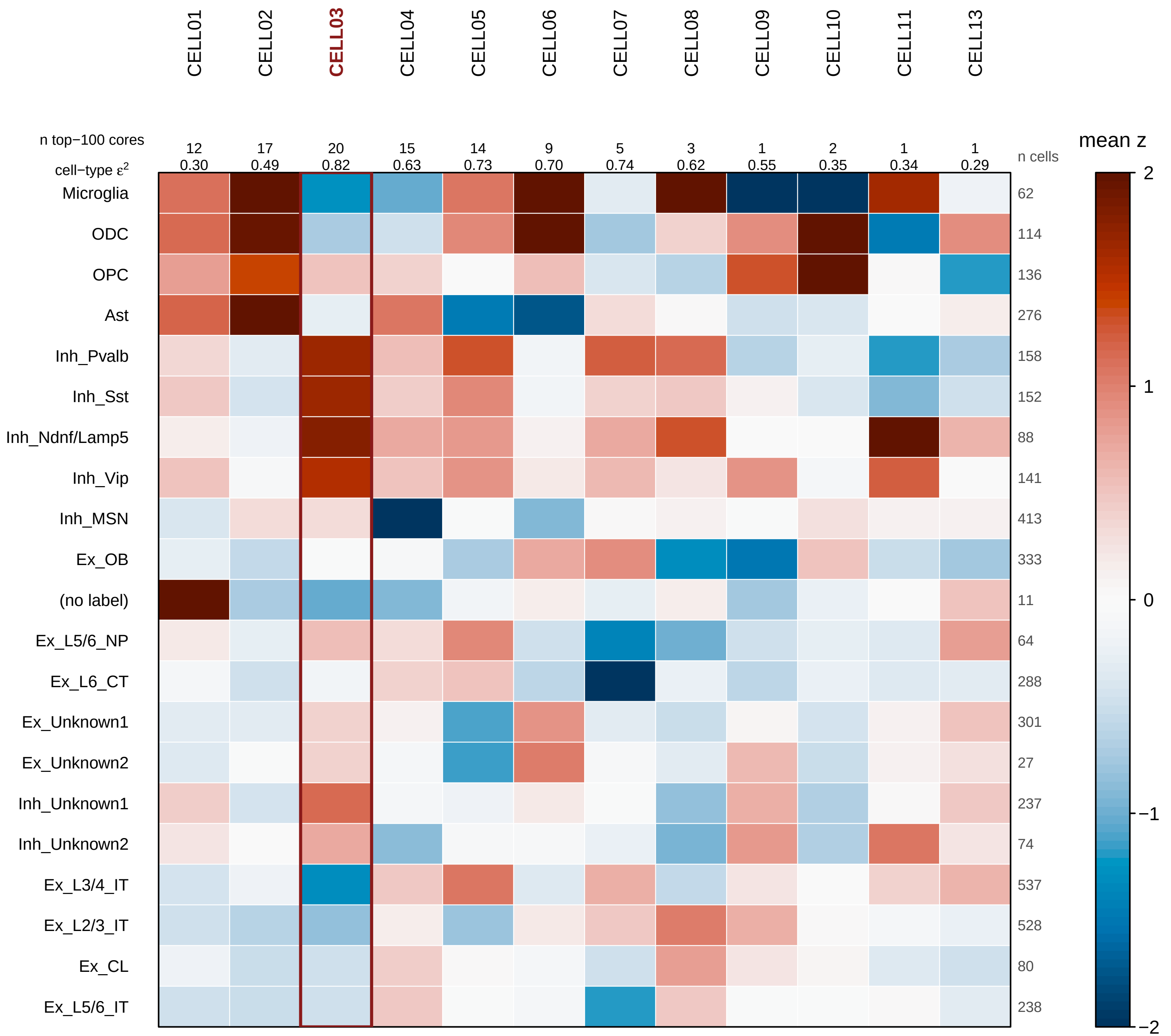

CELL01
CELL02
CELL03
CELL04
CELL05
CELL06
CELL07
CELL08
CELL09
CELL10
CELL11
CELL13
n top−100 cores
12 17 20 15 14 9 5 3 1 2 1 1
cell−type ε²
0.30 0.49 0.82 0.63 0.73 0.70 0.74 0.62 0.55 0.35 0.34 0.29
n cells
Microglia 62
ODC 114
OPC 136
Ast 276
Inh_Pvalb 158
Inh_Sst 152
Inh_Ndnf/Lamp5 88
Inh_Vip 141
Inh_MSN 413
Ex_OB 333
(no label) 11
Ex_L5/6_NP 64
Ex_L6_CT 288
Ex_Unknown1 301
Ex_Unknown2 27
Inh_Unknown1 237
Inh_Unknown2 74
Ex_L3/4_IT 537
Ex_L2/3_IT 528
Ex_CL 80
Ex_L5/6_IT 238
mean z
2
1
0
−1
−2
